# Nature vs nurture? Light availability drives phenotypic plasticity within a reef coral species

**DOI:** 10.1101/2025.09.30.679437

**Authors:** Hugo Ducret, Christopher R. Suchocki, Claire J. Lewis, Heidi Dierssen, Eric J. Hochberg, Daniel W. H. Schar, Marc Kochzius, Isabelle F. George, Robert J. Toonen, Jean-François Flot

## Abstract

Scleractinian corals exhibit wide intra-specific phenotypic variations. However, the extent to which these variations are explained by genotypic variation or phenotypic plasticity remains unclear. To elucidate this question, we devised a replicated experiment in which nine *Montipora capitata* coral colonies were cut into six pieces each, three of which were placed in shaded conditions (with a 73% reduction in light intensity) whereas the other three were kept in natural conditions as a control. After one year, we detected statistically significant morphological differences between corals of the same genotypes grown in different light environments, but no significant change in symbiont community structure. Colonies kept in control conditions exhibited high surface complexity and converged towards more digitate or corymbose morphs compared to shaded corals, which exhibited planar-like surfaces and converged toward laminar and foliose forms. Our data demonstrate the high level of light-driven phenotypic plasticity of *Montipora capitata* and suggest a trade-off between the amount of biomass per space area and the amount of absorption of incident light.

## Introduction

Zooxanthellate corals are sessile marine organisms inhabiting oligotrophic waters. They harbor in their tissues photosynthetic dinoflagellates that contribute to their daily metabolic requirements (Odum *et al*., 1971; Muscatine, 1990; Oakley & Davy, 2018). Because of that, just like plants in tropical forests, corals on a reef compete for light and substrate (Connell, 1978), and light is considered a critical component of their fundamental niche (Hoogenboom & Connolly, 2009). As a result, coral reefs are characterized by diverse assemblages of species (Dornelas *et al*., 2006; Tamir *et al*., 2019; López-Londoño *et al*., 2022), life-history traits and strategies (Madin *et al*., 2014; Zawada *et al*., 2019a), and demographic rates (Dornelas *et al*., 2017).

Light intensity drives several key aspects of coral phenotype. For instance, changes in colony morphology commonly occur between shallow and deep habitats (Chappell, 1980; Muir & Pichon, 2019; Kramer *et al*., 2020). Three main aspects of the morphology of hard corals have been identified by Zawada *et al*. (2019a) and described as their three main axes of morphological variation: surface area complexity, skeletal volume compactness, and the vertical distribution of their volume. Species with planar skeletal surfaces that maximize incoming light per space unit are more frequent in deep, low-light environments (Chappell, 1980; Muir & Pichon, 2019; Kramer *et al*., 2020). Another morphological change increasing light capture is increased top-heaviness, i.e., tabular-like morphologies (Zawada *et al*., 2019b), which reduces the growth and recruitment rates of neighboring colonies by intercepting light (Baird & Hughes, 2000) but results in more fragile colonies (Hughes & Connell, 1999; Madin & Connolly, 2006). To the contrary, species with more complex skeletal surfaces that pack more biomass per space unit (Kruszyński *et al*., 2007; Hoogenboom *et al*., 2008; Muir & Pichon, 2019) are prevalent under high irradiance levels, possibly because compact volumes and rough shapes reduce photodamage through self-shading (Muko *et al*., 2000; Ow & Todd, 2010; López-Londoño *et al*., 2022). These cited examples of phenotypes not only differ in their shapes, but also in their ability to reflect and absorb light at different wavelengths (Joyce & Phinn, 2002; Todd, 2008; Kramer *et al*., 2022). Variations in skeletal features can modulate reflectance at the surface of the coral tissue: for instance, Gomez-Campo *et al*. (2024) showed a light-associated increase in corallite size in *Acropora palmata*, resulting in self-shading. Indeed, these skeletal structures generate a heterogeneous light environment, which decreases the amount of tissue and number of polyps exposed to light. Another process known as multiple scattering, or the diffusion of light in multiple directions by the coral skeleton (Enríquez *et al*., 2005), was shown to increase with the complexity of the surface of coral colonies (Marcelino *et al*., 2013). As a result, more branching increases light absorption for a given amount of photopigments per symbiont cell (Enríquez *et al*., 2005). By contrast, the lower reflectance values of low-light-adapted morphotypes maximize the absorption of incident light (Kaniewska *et al*., 2008; Lesser *et al*., 2021).

Another critical aspect of coral phenotype that is known to change with light habitat is the prevalence of different symbiont types. In the wild, corals can associate with different types of photosymbionts (LaJeunesse *et al*., 2010; Carradec *et al*., 2020), and their distribution is influenced by multiple factors. Among these, depth correlates with intraspecific variations in symbiont community compositions (Warner *et al*., 2006; Frade *et al*., 2008b; Bongaerts *et al*., 2013, 2015). For instance, Frade *et al*. (2008a) found depth-associated clades and shallow-specialist clades of *Symbiodinium* in closely related coral species of the genus *Madracis*. Alternatively, for other coral species, the relative prevalence of symbiont clades varies with depth (Innis *et al*., 2018), with a higher proportion of light and heat tolerant symbionts in shallow habitats (De Souza *et al*., 2022, 2023). This can be explained by differences in concentrations of photopigments in their cells. Depth specialist clades were found to have naturally higher concentrations of the light-harvesting pigments chlorophyll *a* and peridinin (Falkowski, 1981; Frade *et al*., 2008b), which increases light assimilation By contrast, shallow specialists were found to contain higher concentrations of carotenoids involved in the xanthophyll cycle, such as diadinoxanthin (Hochberg *et al*., 2006; Frade *et al*., 2008b). A photoprotective process, commonly known as non-photochemical quenching, (Roth, 2014; Hochberg *et al*., 2006; Shi *et al*., 2018) involves the transformation of diatoxanthin into diadinoxanthin. As this reaction dissipates excess light as heat, it advantages corals found in habitats characterized by high irradiance levels (Roth, 2014).

Although variations in coral phenotype with depth are frequently mentioned in the literature (e.g. Veron, 1995), there is no clear explanation of the factor(s) responsible for these variations. In corals, phenotypic variations are often associated with genetic differentiation (Stefani *et al*., 2011; Schmidt-Roach *et al*., 2014; Smith *et al*., 2017), which can occur either within or between species. Phenotypic plasticity (i.e., morphological differences between genetically indistinguishable individuals because of environmental parameters; West-Eberhard, 1989) has been proposed to be prevalent in many corals, but actual evidence so far concerns mostly micro-skeletal features (Bhagooli, 2003; Todd, 2008; Tambutté *et al*., 2015; Malik *et al*., 2021; Gomez-Campo *et al*., 2024), whereas analyses of colony-scale morphological plasticity have rarely been performed. Similarly, the coral-algal symbiosis can vary through time within individuals, particularly after a bleaching event (Baker *et al*., 2004; De Souza *et al*., 2023), but patterns of symbiosis flexibility are complex and differ across species (Baker, 2003; Baird *et al*., 2007; Ziegler *et al*., 2015). Two main hypotheses can thus be proposed to explain the phenotypic variations observed across environments in corals: i) the observed intraspecific differences are explained by genetic differentiation and environmental filtering (Keddy, 1992), with different genotypes growing at different depths and exhibiting different symbiont community compositions; or ii) the observed differences are the result of phenotypic (Todd, 2008) and symbiont association plasticity (Cunning *et al*., 2015) in response to different environmental conditions. In the latter case, plasticity may be influenced by light intensity and/or by other factors that vary with depth, such as turbidity and water movement (Todd, 2008).

A good model organism to look at the extent to which light intensity can induce plasticity in hard corals is *Montipora capitata*. This species, commonly referred to as the Hawaiian rice coral, is characterized by an unusually high depth range, ranging from one up to one hundred meters below sea level (Veron, 2000). It exhibits a wide spectrum of morphologies, ranging from highly branching to plate-like (Veron, 2000; Padilla-Gamiño *et al*., 2013). Bhagooli (2003) also reported differential growth of tuberculae, or “rice grains”, at the surface of the skeleton depending on its light environment, which they hypothesized to be photoprotective. Similarly, *M. capitata* harbors two main clades of photosymbionts. While mainly dominated by *Cladocopium* spp. (”Clade C”), the proportion of colonies dominated by *Durusdinium glynnii* (”Clade D”), increases in shallow habitats in Kāne‘ohe Bay, Hawai‘i (Innis *et al*., 2018; De Souza *et al*., 2022, 2023).

To dissect whether light-driven plasticity explains the phenotypic variations observed in *Montipora capitata* colonies in the field, we conducted a one-year experiment wherein we grew fragments of identical *Montipora capitata* genotypes in two contrasted light conditions at the same depth. At both the start and the end of the experiment, we measured colony surface area complexity, skeletal volume compactness, distribution of volume vertically in the water column, and symbiont community compositions. We combined these phenotypic traits with estimates of photopigment concentrations, and tested for the effects of light treatment and genotype on the measured traits. We expected to find morphological and symbiont community differences between light conditions.

## Methods

### Experimental design

The experimental design of this study was carried out at the Hawai‘i Institute of Marine Biology, Kāne‘ohe Bay, Hawai‘i. In November 2021, nine *M. capitata* coral colonies were fragmented in six pieces, half of them were shaded by 73% and the other half were grown unshaded (n = 54). Fragmentation was done in a “pie-slice” way, meaning that each resulting fragment contained both peripheral (i.e. younger) and central (older) parts of their mother colony. Each resulting fragment measured approximately 15 cm in diameter at the start of the experiment. As our goal in this study was to cover the full spectrum of traits that *M. capitata* can possibly occupy in the wild, we chose our colonies randomly, without categorizing apriori into “branching” vs. “plating form”. Experimental racks were placed at a depth of 1.5 meters, and top-down photographs of each colony were taken every six months (Fig. 1). HOBO Pendant data loggers (Onset, USA) were placed on each rack to record light intensity over the entire course of the experiment, in five-minutes intervals. Mean daily maxima was around 610 µmol.m^-2^.sec^-1^ in the control treatment, and 90 µmol.m^-2^.sec^-1^ in the shade treatment (Ducret *et al*., 2024).

**Figure 1.**
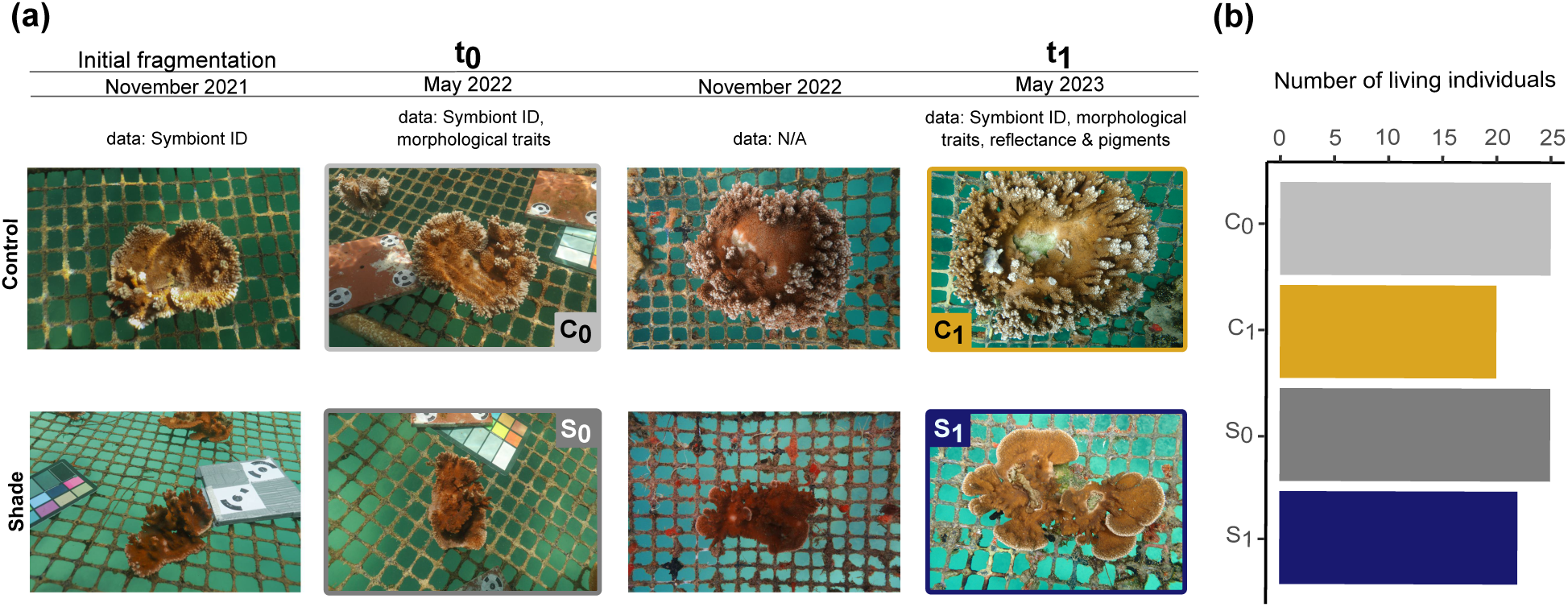
(a) Time-series photographs of one genotype during the one-and-a-half-year period of the experiment in the control (top row) and shade (bottom row) treatments. The meshes in the background display square patterns that are approximately 4×4 cm2. For each time-point, details about the types of data that were collected is available. The t_0_ we refer to in this study corresponds to the series of scans performed six months into the experiment (May 2022), while our ‘t_1_’, corresponds to the scans performed one year after t_0_ (May 2023). Similarly, colored rectangles around photographs and acronyms in the photographs indicate the groups to which these corals belong: light gray = control at t_0_ (C_0_); gold = control after one year (C_1_); dark gray = shade at t_0_; (S_0_); dark purple = shade after one year (S_1_). No data was collected in November 2022, but pictures were taken to monitor coral growth halfway between t_0_ and t_1_. These pictures are shown for illustration purposes. (b) Column plots showing the number of living coral fragments in each aforementioned group.

### Morphological data acquisition

Each coral was 3D-modeled twice using an Artec Space Spider Scanner (Artec 3D, Luxembourg) and the Artec Studio 14 software. Due to technical and logistical constraints, our first round of scans (”t_0_”) was performed six months into the experiment (Fig 1) under both light conditions. However, given the large size of our coral fragments, new growth was relatively minimal at this point, and no trait showed a significant difference between control and treatment t_0_ groups (Supplementary Table S2). The second round of scans was performed one year later (”t_1_”).

Colonies were taken out of the water for scanning, and during scan processing, they were placed into a flow-through water table to avoid air exposure. Four fragments died between the initial fragmentation (November 2021) and the first round of scans (t_0_, May 2022) and were excluded from the analysis. Scanning was completed using the real-time fusion mode for all fragments until all surfaces were covered. Once scanning was complete, we used the global registration algorithm to compile all scans together before fusion (Reichert *et al*., 2016). The sharp fusion algorithm was used to compute one 3D model from all previously-generated scans for each colony. We subsequently applied the “small objects filter” algorithm to get rid of artifacts generated by shading or reflection. The final models were subsequently exported as .obj files for downstream analyses.

### Morphological data analysis

We aimed to compute quantitative, biologically relevant morphological traits to capture the above-mentioned axes of variation at the colony scale (Zawada *et al*., 2019a). To do so, we used the `habtools R` package (Schiettekatte *et al*., 2025) on the previously-generated 3D models. Prior to computing morphological traits, the resolution of the models was assessed using the `Rvcg::vcgMeshres` command in RStudio. We first extracted the size descriptors, volume and surface area, from the polygon meshes. Then we computed predictors of surface complexity, i.e. the surface area-to-volume ratio (SA:V) and rugosity (ratio of surface area to planar area). We then computed convexity (the ratio of the volume of an object to the volume of its convex hull, which is a measure of volume compactness) and top-heaviness (a measure of the distribution of the volume of an object along its vertical axis; Schiettekatte *et al*., 2025). Details about the mathematical formula for each variable are available in the Supplementary Table S1.

### Optical property measurements

Coral reflectance was measured in May 2023 (Fig. 1) using a hand-built dive spectrometer (Russell & Dierssen, 2023). The instrument consists of multichannel Jaz spectrometers (Ocean Optics) inside a customized underwater housing. Sampling was performed using a 600 µm fiber optic probe pointing at the targetted sample, and a 20% gray-calibrated Spectralon plaque (Labsphere) was used to standardize our measurements (Russell & Dierssen, 2023). The reflectance of the calibration plaque was measured before measuring the reflectance of each colony, to control for fluctuating light conditions and to assign a quality metric to each measurement. If light conditions changed dramatically during the course of sampling and created large shifts in the spectrum viewed by the diver (e.g., changing cloud cover), sampling was truncated and noted in the metadata. Reflectance was calculated as the ratio of radiance from the coral relative to that from the calibrated 20% gray Spectralon reflectance target, which was held horizontally and at the same depth as the coral. Because there is a high heterogeneity of light gradients and optical microniches in coral colonies (Wangpraseurt *et al*., 2012), reflectance was analyzed on 4 growing edges of fragments and on 5 samples near the center of fragments which led to n = 9 samples per coral fragment. Due to large uncertainty arising from wave focusing or “caustics” (a phenomenon caused by light hitting the irregular sea surface at different angles), the direct beam of light had to be shielded over the target when making measurements. Therefore, the target and plaque were shaded with one cupped hand placed vertically in front of the targets in the same configuration (Hochberg *et al*., 2003).

As spectral reflectance patterns of corals are largely driven by the pigments of photosynthetic symbionts within the coral host, the relative concentration of photopigments can be related to the reflectance values (Hochberg *et al*., 2006). We thus estimated the concentrations of two photopigments, namely chlorophyll *a* and diadinoxanthin, based on the obtained reflectance values using a partial least-squares regression (PLSR) following the model and methods described in Hochberg *et al*. (2006). However, we used new coefficients for the PLSR, which are available in the online supplementary material (modified from Hochberg *et al*. (2006), Supplementary data file “pigments_PLSR_coefficients.xlsx”). Differences in pigment concentrations between treatments and locations on the coral colony were assessed using two-way Analyses Of Variance (ANOVAs), and difference in reflectance spectra were assessed using Mann-Whitney tests.

### Zooxantellae typing

In order to investigate whether light intensity induces changes in symbiont community structure (Innis *et al*., 2018; De Souza *et al*., 2022), vertical branch tip samples of *M. capitata* corals were collected after fragmentation and prior to treatment application (November 2021), after six months (May 2022, during the first round of scans), and after a year and a half of experiment (May 2023, during the second round of scans, Fig. 1). All samples were collected at solar noon, and stored in DNA/RNA shield buffer (Zymo Research, USA) at −20°C immediately after collection. Prior to performing DNA extraction, samples were ground in a bead-beater (Precellys instruments) at the following conditions: speed: 6500rpm, time: 30sec, pause time: 60sec, cycles: 3 (Belser *et al*., 2023), and aliquoted for downstream analyses. Hereafter, 10µl of Proteinase K (20 mg.ml^-1^) were added to the mix, and samples were incubated at 55°C for 30sec. DNA extraction was performed using Macherey-Nagel’s Genomic DNA from tissue extraction kit, following the manufacturer’s instructions.

The Internal Transcribed Spacer 2 (ITS2) region of the rDNA of zooxanthellae was amplified using a two-step PCR as detailed in Collard *et al*. (2025). For the inner primers we used the primer pair developed by Hume *et al*. (2018), to which we added either a forward tail sequence, or a reverse tail sequence, retrieved from the Oxford Nanopore Technologies (ONT) barcoding kits. The resulting inner primer sequences were as follow; Forward primer: 5’-TTTCTGTTGGTGCTGATATTGCGAATTGC-AGAACTCCGTGAACC-3’, Reverse primer: 5’-ACTTGCCTGTCGCTCTATCTTC-CGGGTTCWCTTGTYTGACTTCATGC-3’. The 25µL PCR mix consisted of 12.5µL of PhireTaq Plant Master Mix 2X (ThermoScientific), 1µL of inner primers at 10µM, 0.5µL of DNA template, and volumes were adjusted with Nuclease-Free water. PCR conditions for the first step were as follow: Initial denaturation at 98°C for 4mins, then 15 cycles of denaturation at 98°C for 8sec, annealing at 56°C for 8sec, and extension at 72°C for 10sec. PCR strips were then taken out of the thermocycler, and a unique barcode pair (Forward-Reverse) was added to each well. The outer primer set consisted in the same tail sequences for binding, and a unique barcode sequence, retrieved from the ONT native barcoding kits (Collard *et al*., 2025). This allowed for pooling multiple samples into one ONT sequencing run. PCRs were then homogenized and re-incubated in the thermocyclers. The second step of PCR consisted in 25 cycles of denaturation at 98°C for 8sec, annealing at 56°C for 8sec, and extension at 72°C for 10sec. PCR products were checked on agarose gel and pooled in equimolar quantities to constitute the input of the sequencing library (Carradec *et al*., 2020). Sequencing libraries were prepared for R10.4.1 flow cells ran on the Mk1C MinION device, using the Ligation Sequencing kit (ref: SQK-LSK 114) from the Oxford Nanopore Technologies (Carradec *et al*., 2020). The SpotON flow cell was loaded according to the manufacturer’s instructions, and each sequencing run was performed for 24 hours.

Basecalling was performed using the `guppy_basecaller` function of `guppy v6.3.8`. The obtained reads were demultiplexed using the `guppy_barcoder function` of `guppy`. We then used the command-line pipeline developed by Carradec *et al*. (2020) to analyze the raw ONT data. Reads were mapped to an ITS2 reference database containing 432 sequences of Symbiodiniaceae (Arif *et al*., 2014) using `minimap2` with “map-ont” as options. The reference sequences covered with a minimum of 1% of all sequences were kept for a second round of read-mapping. This second round was performed in order to aggregate reads potentially mis-assigned during the first round of read-mapping (Carradec *et al*., 2020). We then used SAMtools and BCFtools to reconstruct consensus sequences (Danecek *et al*., 2021), and samples were rarefied at a depth of 1000 sequences per individual using the Rarefy function of the *vegan* R package (Oksanen *et al*., 2024).

### Statistical analyses

All computed morphological traits were gathered in a matrix used for subsequent analyses. Individuals were grouped as follow; control at t_0_ (hereafter referred to as “C_0_” group), shade at t_0_ (“S_0_” group), control at t_1_ (“C_1_” group) or shade at t_1_ (“S_1_” group, Fig. 1). We first aimed at assessing how light intensity impacted the measured morphological traits individually. To make sure that no difference was already present at t_0_, we compared the values of each trait between C_0_ and S_0_ using one-way Analyses of Variance (ANOVAs). Subsequently, we measured the log-ratio (lr) or each variable for each individual. Individuals that were not scanned at both t_0_ and t_1_ due to mortality were excluded, which led to n = 42 observations. We then built a linear mixed model with the `lmer()` function of the `lme4` package in RStudio (Bates, 2010), with genotype as a random effect and treatment as a fixed effect, and tested for the significance of treatment on each morphological trait. The same approach was used with the `glmer()` function of the same package to test for the effect of treatment on mortality, which was treated as binary.

We then used the dudi.pca() function from the ade4 package in RStudio (Chessel *et al*., 1995) to plot a principal component analysis (PCA). This was made in order to look at morphospace overlap between these four groups, and identify how morphological traits changed collectively during the year of experiment. For this multivariate analysis, variables were log-transformed to reduce the influence of variable scale on the ordination plot. Homogeneity of variances of the four groups (C_0_, S_0_, C_1_, S_1_) was assessed using a Permutational Multivariate Analysis of Dispersion (PERMDIST) performed via the betadisper() function of the vegan R package (Oksanen *et al*., 2024). We then tested for significant morphological differences between groups and between genotypes using Permutational Analyses of Variance (PERMANOVAs; adonis2() function, vegan R package Oksanen *et al*., 2024). Similarly, effects of treatment and genotypes on symbiont communities were also assessed through PERMANOVAs. Additionally, we looked at autocorrelation of untransformed variables using Pearson’s r correlation coefficients, to see how variables covary. Lastly, we computed the Euclidean distances between paired individuals and the angles of orientation of the corresponding vectors in our PCA of morphological traits. This was performed to quantify the amount of distance traveled and its orientation between paired PCA points. For each pair of points, distance and angle were calculated as 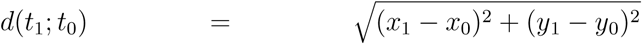, and *θ*(*t*_1_; *t*_0_) = *arctan*((*x*_1_ *− x*_0_)*/*(*y*_1_ *− y*_0_)), respectively, and were compared across treatments and genotypes. Lastly, the Inkscape program was used to add photographs of coral colonies and illustrations of the different shape variables on the generated figures (**?**).

## Results

### Models and morphological traits

3D models were successfully generated and are available under the following link: https://mycloud.ulb.be/index.php/s/57oYmHzK5Y42ZR6. Additionally, time-series photographs of the studied coral colonies are available in supplementary material (Supplementary Figures S5 to S13). An example is provided on Fig.1 for illustration purposes. Overall, few fragments died during the experiment (Fig. 1b), with no significant influence of treatment or genotype on the latter (mixed model, p-values > 0.05). No trait showed a significant difference between C_0_ and S_0_ groups (one-way ANOVAs, p-values > 0.05, Supplementary Table S2). Surface area, SA:V and top-heaviness significantly covaried with volume, while rugosity and convexity did not (Supplementary Fig. S2).

Log-ratios of volume, surface area, SV ratio and top-heaviness were significantly different from zero (Supplementary Tables S3, S4; mixed models, p-values_intercept_ < 0.05), which shows that these variables significantly changed after one year. Light intensity showed a significant effect on all measured traits (Fig. 2a, Supplementary Table S4, p.values < 0.05). Colonies in the S_1_ group showed higher SA:V ratios, while colonies in the C_1_ group were characterized by a higher values in volume, surface area, rugosity and convexity (Fig. 2). The increase in top-heaviness is also significantly higher in the C_1_ group, but its corresponding p-value is just below the significance threshold (Table S4, p-value = 0.47). As such, the statistical power is rather low, and reliable conclusions on the significant increase of this variable with light is unlikely.

**Figure 2.**
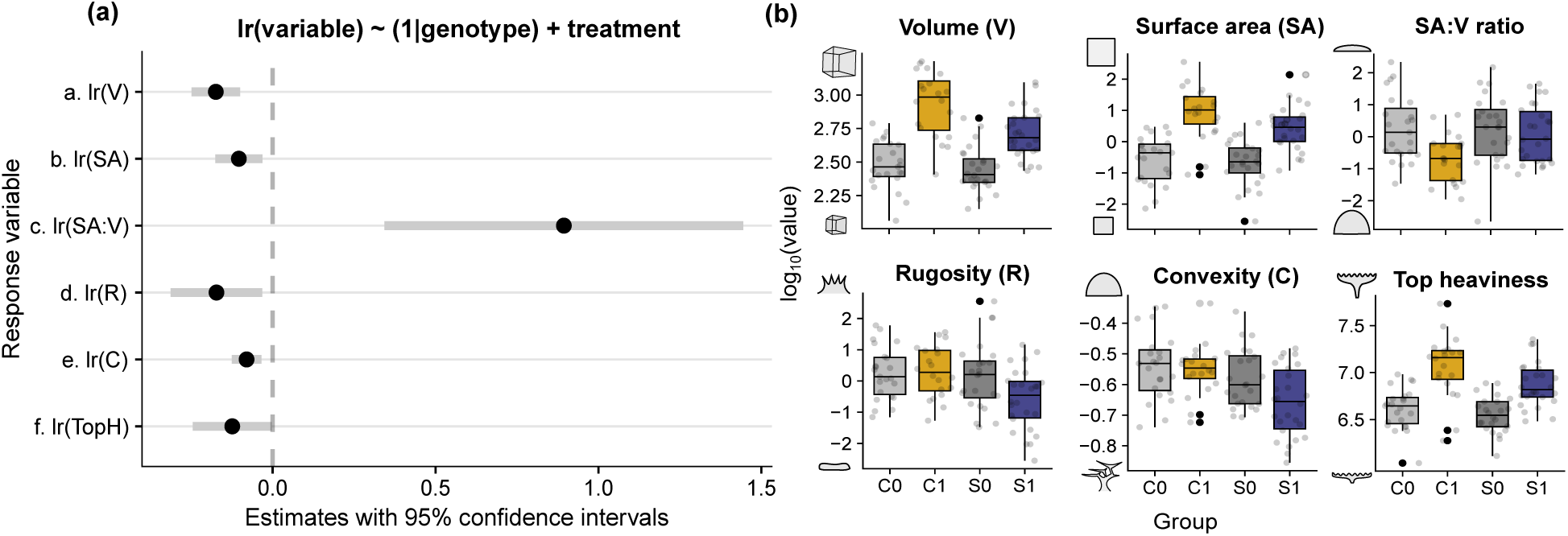
(a) Results of our linear mixed models, showing the effect size of light treatment on the log-ratio (lr) of each morphological trait between t_1_ and t_0_. Black dots represent variables for which treatment has a significant effect (here, effect is significant for all variables). Gray bars depict 95% confidence intervals. (b) Boxplots showing the distribution of log-transformed values of each measured trait across the four groups (C_0_ = control at t_0_; C_1_ = control after one year; S_0_ = shade at t_0_; S_1_ = shade after one year).

### Reflectance and photopigments

We compared the reflectance of growing edges and bodies of corals grown under ambient light or reduced light at t_1_. We found a significant difference in reflectance among treatments, as well as between bodies and growing edges of the same treatment (Fig. 3a; Mann-Whitney, p-value < 0.05). Typically, mean reflectance values were significantly lower in shaded corals compared to control corals (Kruskal-Wallis, p-value < 0.05). Additionally, in both treatments, mean reflectance values were significantly higher at the growing edges than in center locations (Kruskal-Wallis, p-value < 0.05).

**Figure 3.**
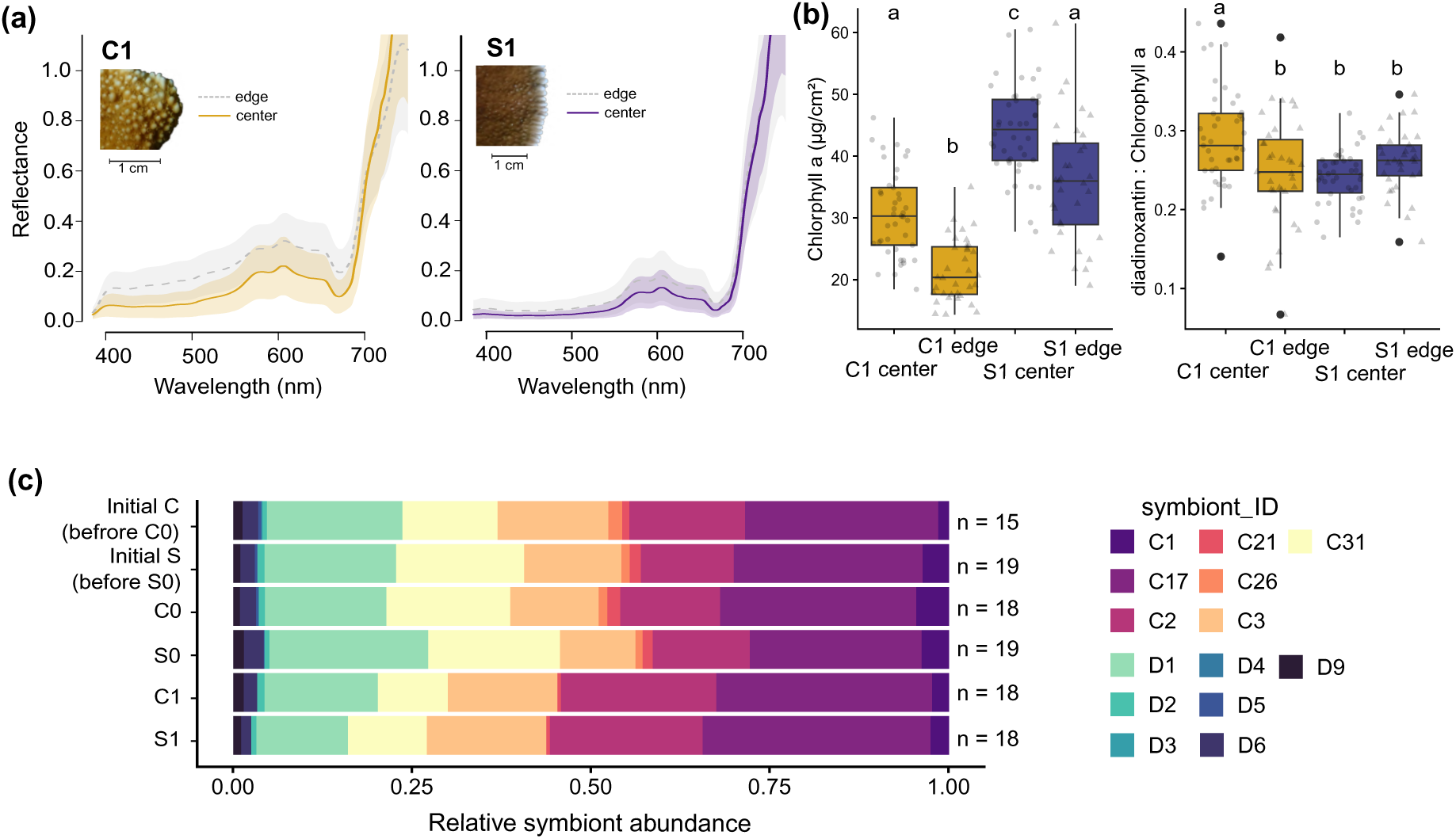
(a) Reflectance of C_1_ and S_1_ groups for the 400-700nm wavelength range. Sold lines represent mean reflectance values measured at center locations, while dashed lines represent values that were measured at edge locations. Shaded areas represent 95% confidence intervals. Additional close-up photographs of colony edges from both treatments were added to emphasize the microscopic morphological variations of *M. capitata*. (b) Chlorophyll *a* concentration (µg.cm^-2^) and diadinoxanthin:chlorophyll *a* ratio derived from the reflectance of control and shaded corals, measured at both center (n = 45 in each group) and edge (n = 36 in each group) locations. Letters above the boxplots denote significance levels. (c) Stacked column plots of symbiont clade prevalence, with means of relative abundances from samples grouped by treatment and time-point.

We estimated significantly higher chlorophyll *a* concentrations in shaded colonies compared to control colonies, as well as in centers compared to growing edges (Fig. 3b; two-way ANOVAs, p-values < 0.01). The diadinoxanthin:chlorophyll *a* ratio was significantly higher at the center of control colonies compared to all other locations and treatments (Kruskal-wallis, p-value < 0.01). However, no statistical differences were detected between the values obtained from the edges of the control colonies and those of the shaded colonies at both edges and centers (Fig. 2b; two-way ANOVAs, p-value > 0.05).

### Symbiont community data

A total of 412 509 symbiont sequences were generated from our *M. capitata* coral samples. After demultiplexing, quality control, and read-mapping this resulted in an average of 3702 sequences per individual. Samples with less than 1000 sequences were excluded from the analysis. Overall, we identified 14 distinct Symbiodiniaceae clades (Fig. 3c), with 7 belonging to the genus *Cladocopium* (Clade C), and 7 to the genus *Durusdinium* (Clade D). C17 was the most abundant Symbiodiniaceae from the genus *Cladocopium* and accounted for about 24% of total hits, while D1 was the most abundant Symbiodiniaceae from the genus *Durusdinium* and represented 23% of the total abundance. We did not find any significant effect of light treatment, time, or a combination of both (PERMANOVAs, p-values > 0.05) on the symbiont community composition. However, we found a genotypic effect on the latter (PERMANOVA, p-value < 0.01); two genotypes were *Durusdinium*-dominated, while the seven others were *Cladocopium*-dominated, and this prevalence did not change over time (Supplementary Fig. S1). Lastly, we did not see any significant effect of symbiont clade dominance on morphospace overlap (PERMANOVA, p-value > 0.05).

### Principal Component Analysis

We used a principal components analysis (PCA) to look at morphospace overlap between each group based on the measured traits. Overall, 81.8% of the total variance was explained by the two first dimensions (Fig. 4). The first principal component (PC1) explained 51.4% of the total variation with surface area, top heaviness, and volume contributing most to this component. The second principal component (PC2) explained 30.4% of the variation, with convexity, SA:V ratio, and rugosity contributing most to this component. The PERMDIST test revealed that average distances from spatial median ranged from 1.336 (C_1_) to 1.485 (S_0_), so we considered the intra-group variances to be homogeneous. At t_0_, we found no significant effect of treatment on coral morphology, suggesting no group effect within the dataset at the start of the experiment (PERMANOVA C_0_-S_0_; p-value > 0.05). After one year of growth, we found a significant change in the morphospaces of the control corals (PERMANOVA C_0_-C_1_; p-value < 0.01), and of the shaded corals (PERMANOVA S_0_-S_1_, p-value < 0.01). Additionally, when assessing differences between treatment groups C_1_ and S_1_, we found a significant effect of light intensity on coral morphology after one year of growth (PERMANOVA; p-value < 0.01). The effect of genotype on coral morphology was also significant (Supplementary Fig. S4; PERMANOVA, p-value < 0.01). Mainly, two genotypes (”B” and “H”) were significantly distinct from the others (Supplementary Table S5).

**Figure 4.**
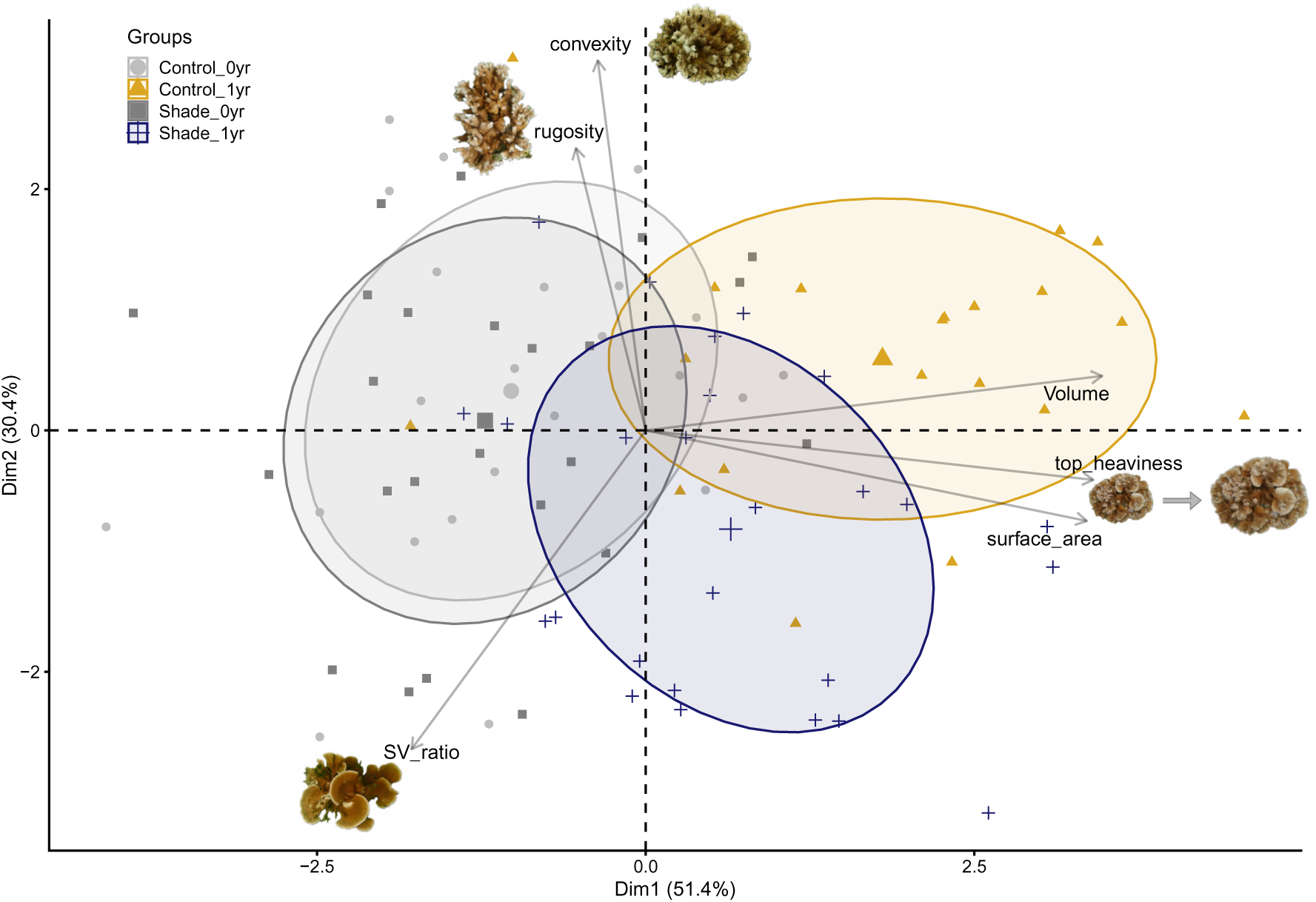
Principal component analysis (PCA) of morphological traits, with individuals grouped by time and treatment, and ellipses depicting 95% confidence intervals. The first principal component (PC1) broadly captures growth through time, while the second (PC2) captures a trade-off between surface area complexity and planarity. Images of *M. capitata* colonies are of the coral specimens that occupy extremes of shape variables.

### Distance and angles between paired PCA points

For our dataset of morphological traits, we calculated the Euclidean distance across paired PCA points, and the angles inclination of the corresponding vectors (Fig. 5a). We did not compute these metrics on our symbiont community dataset as no change was detected between paired individuals through time or among treatments (Supplementary Figure S1). We found no significant effect of treatment on the distance traveled (Fig. 5b; linear mixed model, p-value > 0.05), but the angles of inclination of the vectors differed between control and shaded individuals (Fig. 5c; linear mixed model, p-value < 0.05). In addition, there was no significant effect of coral genotype on distance traveled or angle of inclination of vectors (p-values > 0.05).

**Figure 5.**
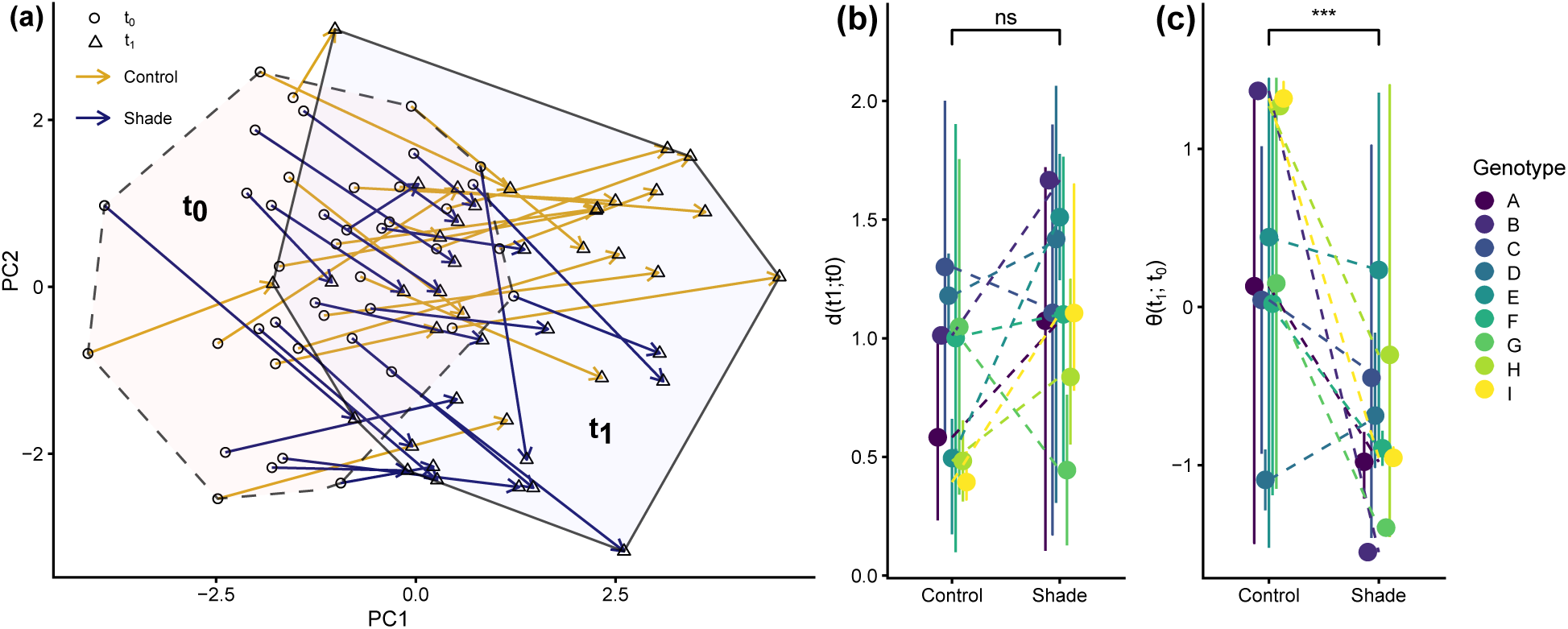
(a) Distance traveled between paired PCA points. Circles represent individuals scanned at t_0_, while triangles denote individuals scanned at t_1_. Similarly, the dashed polynom represents the morphospace of individuals scanned at t_0_, while the solid polynom represents the morphospace of individuals scanned at t_1_. Arrows are color-coded by treatment. (b) Euclidean distances between paired individuals for each genotype in each treatment. (c) Angles of inclination of vectors between paired PCA points for each genotype in each treatment. Dashed lines allow to visualize the differences in mean traveled distance or angle of inclination between identical genotypes growing in different groups or treatments. The displayed significance levels correspond to the significance of the “treatment” factor in our mixed model with genotype included as a random effect.

## Discussion

In this study, we show evidence of light-induced morphological plasticity in *Montipora capitata* corals, which exhibited distinct morphotypes in response to their light environment. Control corals tend toward more digitate or corymbose forms, while shaded corals tend toward laminar and foliose forms (Fig. 1, Fig. 4). However, we did not detect any change associated with light intensity in symbiont type prevalence (Fig. 3c), which variations were more associated to a genotypic effect. The different axes of phenotypic variation are discussed below.

### There is a trade-off between biomass per surface area and absorption of incident light

The higher rugosity and lower SA:V ratio we observed in the C_1_ group compared to the S_1_ group reflect differential variations in surface complexity that have been hypothesized as a functional trade-off between biomass packing and decreased intra-colony competition for resources (Hoogenboom *et al*., 2008; Wangpraseurt *et al*., 2012; Zawada *et al*., 2019a). Increased surface complexity and reflectance in the control treatment may allow for maximized light assimilation via enhanced multiple scattering, thus minimizing investment in light-harvesting pigments (Fig. 3b, Marcelino *et al*., 2013). By contrast, the planar-like morphologies of shaded colonies minimize self-shading and generate a more homogeneous light habitat (Fig. 1, Fig. 2), with, supposedly, optimized absorption of incident light via higher concentrations of chlorophyll *a* (Hochberg *et al*., 2006), particularly at central locations (Fig. 2b). This shows that the observed morphological variations are driven by constraints in light availability (Vermeij & Bak, 2002; Hoogenboom *et al*., 2008; Kramer *et al*., 2020), and provides physiological insight to explain why maximum coral diversity is achieved at intermediate depths, i.e., in environments with moderate irradiance levels (López-Londoño *et al*., 2022). In addition, these morphologies clearly resonate with the differences in the growth patterns seen in shallow and mesophotic coral colonies (Lesser *et al*., 2018; Muir & Pichon, 2019) and with the morphological variations one can observe in the reefs of Kāne‘ohe Bay for that species (Jokiel, 1991).

### Volume compactness increases photoprotection

The higher diadoxanthin:chlorophyll *a* ratios in C_1_ colonies suggests that these corals may be experiencing photodamage (Roth, 2014; Shi *et al*., 2018), but have a better ability to release excess energy through non-photochemical quenching, (Hochberg *et al*., 2006). Additionally, the higher convexity values observed in the C_1_ group compared to the S_1_ group denote higher volume compactness to cope with high irradiance levels. This pattern is consistent with what has been documented in arborescent coral species (Todd, 2008; Kruszyński *et al*., 2007), even though the degree to which *M. capitata* forms branches is highly variable (Fig. 1). These mechanisms provide corals with a competitive advantage under high light conditions Frade *et al*. (2008a).

In addition, the high rugosity values seen in C_1_ colonies compared to S_1_ colonies may be partly due to the higher formation of tuberculae at the surface of their skeleton (Fig. 3a), which create more heterogeneous surfaces. Indeed, *Montipora capitata* is known to exhibit a significantly higher rate of tuberculae under high light conditions (Bhagooli, 2003). These upright structures are typical of *Montipora* species. They were defined as projections of coenosteum on the surface of the skeleton that are more than a corallite in width (Veron, 2000; Bhagooli, 2003). Such micro-skeletal features are usually key in modulating light incidence angle, self-shading, inter-reflections, and a generally complex interaction with the light environment (Joyce & Phinn, 2002; Ow & Todd, 2010; Wangpraseurt *et al*., 2012; Lesser *et al*., 2021; Rocha *et al*., 2014; Gomez-Campo *et al*., 2024). As such, higher surface rugosity due to growth of white, symbiont-free tuberculae may explain the significantly higher reflectance ranges obtained in control corals (Fig. 2c). This would confirm the self-shading hypothesis previously described by Bhagooli (2003). However, we highlight that our data focuses on colony-scale metrics. As a result, the detailed, micro-scale morphology of the tuberculae cannot be explicitly quantified. To do so, we recommend examining morphology at multiple scales for the future, by -for example-using fragments with smaller surface areas.

### *M. capitata*’s top-heaviness does not increase at low light

We hypothesized that morphological changes for light capture could increase colony height and top-heaviness under shade, making shaded colonies more mechanically vulnerable (Madin & Connolly, 2006). However, in our experiment, we did not find any significant increase in top-heaviness in the S_1_ group compared to the C_1_ group (Fig. 2a). This may be explained by the fact that phenoptypic plasticity is restrained by species-specific (i.e. genetic) boundaries. *M. capitata* can harbor a wide variety of morphologies, but these do not include top-heavy ones such as tabular morphotypes (Veron, 2000; Zawada *et al*., 2019a). Corals with tabular morphologies are often morph specialists, which is probably the result of a conserved competitive advantage (Baird & Hughes, 2000), with an associated specific demographic strategy (Álvarez-Noriega *et al*., 2016; Dornelas *et al*., 2017). Morphological variations can occur in tabular corals (Ramírez-Portilla *et al*., 2022), but they are not as radical as the ones observed here. However, we found that top-heaviness increased with colony size (Supplementary Fig. S2), suggesting that for this species, mechanical vulnerability increases with colony size, as observed by Madin *et al*. (2014).

### Morphological plasticity occurs without changes in symbiont community

We show that, for each genotype, symbiont community structure did not change over the course of the experiment (Fig. 3c, Supplementary Fig. S1). For *M. capitata*, acclimation to different levels of light intensity may primarily be done via morphological plasticity and photopigment regulation, rather than via a flexible symbiont association (Fig. 3b, Roth, 2014, Ziegler *et al*., 2015). Alternatively, symbiont species composition may be more sensitive to the spectral composition of the light field (i.e., the color of the water) rather than to the intensity of light (Frade *et al*., 2008b). The pigments in coral symbionts primarily absorb blue and red light rather than green light and may be sensitive to the spectral shape of light in the water column. Prior studies have found that the same species of coral can have different symbiont types depending on the location of the reef and the clarity or optical properties of the water (LaJeunesse *et al*., 2010; Russell *et al*., 2019). Here, water clarity was consistent across treatments and shading primarily changed the magnitude of the light field equally across the spectrum. Hence, the relative spectral composition of light was similar for both treatments. This may also explain why Bhagooli (2003) did not find any changes in symbiont type prevalence associated to light intensity in his study on *M. capitata* in Kāne‘ohe Bay, as opposed to studies using depth as the explicative variable rather than light intensity (Innis *et al*., 2018; Wall *et al*., 2020; De Souza *et al*., 2022, 2023). Future studies could consider changing the spectral composition of light between treatments to mimic differences in water clarity and depth to determine if this is a greater determinant of symbiont composition.

### Shaded corals and control ones exhibit divergent growths

These treatments induced growth toward significantly different morphotypes. However, because initial coral fragments were already relatively large in size (12-15cm in diameter), with disparate morphologies (Fig. 1), the morphospace of all individuals at t_0_ was also relatively wide (Fig. 5a, for some contrast see Brambilla *et al*., 2021). As seen above, what changed most between corals of the C_1_ group and those of the S_1_ group is not the size of their morphospaces (Fig. 4), nor the magnitude of the changes between t_0_ and t_1_ (Fig. 5a,b). Instead, it is *θ*, the direction of the changes (Fig. 5c). Vectors of paired points of shaded individuals were more inclined than those of control individuals. The effect of genotype on *θ* was not significant, which may be attributed to a strong intra-genotypic variability in *θ* (Fig. 5c). Nevertheless, the magnitude of the difference in *θ* between treatments differed across genotypes. For instance, the difference in *θ* between control and shade for genotype “I” is larger than for genotypes “E” and “C”, which suggests that the former is phenotypically very plastic. To the contrary, *θ* was higher in the shade for genotype “D”, probably because this genotype displayed non-convex shapes regardless of light conditions (Supplementary Figure S8). This, along with the significant effect of genotype on coral morphology (Supplementary Fig. S4), suggests that the extent to which corals can express phenotypic plasticity varies among individuals. This resonates with the commonly observed genotype-by-environment interaction in such experiments with corals (Ow & Todd, 2010; Drury & Lirman, 2021; Brambilla *et al*., 2021).

## Data and code availability

The R codes used in this study are available on the Github repository through the following link: https://github.com/montiporacapitata/lightinducedphenotypicplasticity/tree/main. The generated 3D models are also available via the following link https://mycloud.ulb.be/index.php/s/57oYmHzK5Y42ZR6 (a Zenodo archive with a permanent DOI will be generated and linked here upon acceptance of the manuscript).

## Acknowledgements

We are grateful to the ‘āina on which this work took place, the ahupua‘a of He‘eia in the moku of Ko‘olaupoko, and acknowledge this as a part of a larger territory recognized by Indigenous Hawaiians as their ancestral grandmother, Papahānaumoku. We acknowledge the generations of Indigenous Hawaiians and their knowledge systems that shaped Hawai‘i in sustainable ways that allow this work to be possible. We hope to honor this relationship by recognizing its foundational importance in this work. This project was funded by the Belgian Fonds de la Recherche Scientifique (F.R.S - Le FNRS) via an “ASP” PhD fellowship (1.A.678.22F) as well as through multiple travel grants to H. Ducret; the COral Reef Airborne Laboratory (CORAL) project of the National Aeronautical and Space Administration and the Ocean Biology and Biogeochemistry program (NNX16AB05G) and National Aeronautics and Space Administration Biological Diversity and Ocean Biology funding (NNX15AC32G) to H. Dierssen; the NASA Grant NTLHJXM55KZ6 to E. Hochberg; the Fonds d’Encouragement à la Recherche (ULB 2018), the Crédit de Recherche “CoGeS” (J.0077.25) and the Projet de recherches “NiGeS” (T.0078.23) to J-F. Flot. We thank Mr. Laurent Grumiau and Mrs. Florence Rodriguez Gaudray for their assist with the laboratiry work. We are thankful to Mr. Tristan Permentier for his help preparing the time-serie photographs, to Dr. Roland Faure for his help with the data analysis, and to Dr. Joshua Madin for his insightful comments on this manuscript.

