## Supplementary Information for "Nature vs nurture? Light availability drives phenotypic plasticity within a reef coral species" (from another version of this preprint: version 1, 2025-10-01)

### Supplementary material

**Table S1.** Description of the computed morphological traits, with their respective axis of variation and formula. Mathematical explanations were retrieved from Zawada *et al.* (2019b).

| Variable | Morphological axis | Explanation | Formula |
|---|---|---|---|
| Volume | Size | The total volume of the object | |
| Surface area | Size | The total surface area of the object | |
| Surface area to volume ratio | Surface complexity | Ratio of surface area volume | $SA/V$ |
| Rugosity | Surface complexity | Ratio of 2D surface area (SA) and 3D surface area ($A_g$) | $R = SA/A_g$ |
| Convexity | Volume compactness | Ratio of the volume of the object ($O_{VOL}$) and the volume of the convex hull around the object ($C_{VOL}$) | $C = O_{VOL}/C_{VOL}$ |
| First moment of surface area | Top heaviness | The total surface area of the object ($O_{SA}$) multiplied by vertical distance from the objects lowest point (H). | $T = \int_{H=0}^{H_{max}} O_{SA} H$ |

**Table S2.** One-way analyses of variance for each variable between $C_0$ and $S_0$ groups.

| Variable | p |
|---|---|
| Volume | 0.496 |
| Surface area | 0.668 |
| SV ratio | 0.935 |
| Rugosity | 0.741 |
| Convexity | 0.186 |
| Top-heaviness | 0.288 |

**Table S3.** Coefficient estimates and confidence intervals for the linear mixed models. Asterisks denote significance levels ($p < 0.05$).

| Dependent variable | lr_volume | lr_SA | lr_SV_ratio | lr_rugosity | lr_convexity | lr_toph |
|---|---|---|---|---|---|---|
| Treatment | -0.174* | -0.104* | 0.894* | -0.172* | -0.080* | -0.084 |
| sd(Treatment) | (-0.246,-0.103) | (-0.173,-0.034) | (0.364,1.424) | (-0.308,-0.037) | (-0.123,-0.036) | (-0.196,0.027) |
| Intercept | 0.449* | 0.349* | -0.467* | 0.002 | -0.017 | 0.416* |
| sd(Intercept) | (0.375,0.522) | (0.286,0.411) | (-0.875,-0.059) | (-0.098,0.101) | (-0.049,0.014) | (0.330,0.503) |
| Observations | 42 | 42 | 42 | 42 | 42 | 42 |
| Akaike Inf. Crit. (AIC) | -34.812 | -40.053 | 118.645 | 8.15 | -83.033 | -5.978 |

**Table S4.** Corresponding p-values of the mixed model coefficients described in Table S3. Asterisks denote significance levels ($* \leq 0.05; ** \leq 0.01; *** \leq 0.001; **** \leq 0.0001$)

| Variable | p(Intercept) | signif | p(treatment) | signif |
|---|---|---|---|---|
| lrV | 2.41E-13 | *** | 3.79E-05 | *** |
| lrSA | 2.33E-12 | *** | 0.006 | ** |
| lrSAV | 0.031 | * | 0.002 | ** |
| lrR | 0.974 | ns | 0.017 | * |
| lrC | 0.296 | ns | 0.001 | *** |
| lrTH | 9.92E-12 | *** | 0.047 | * |

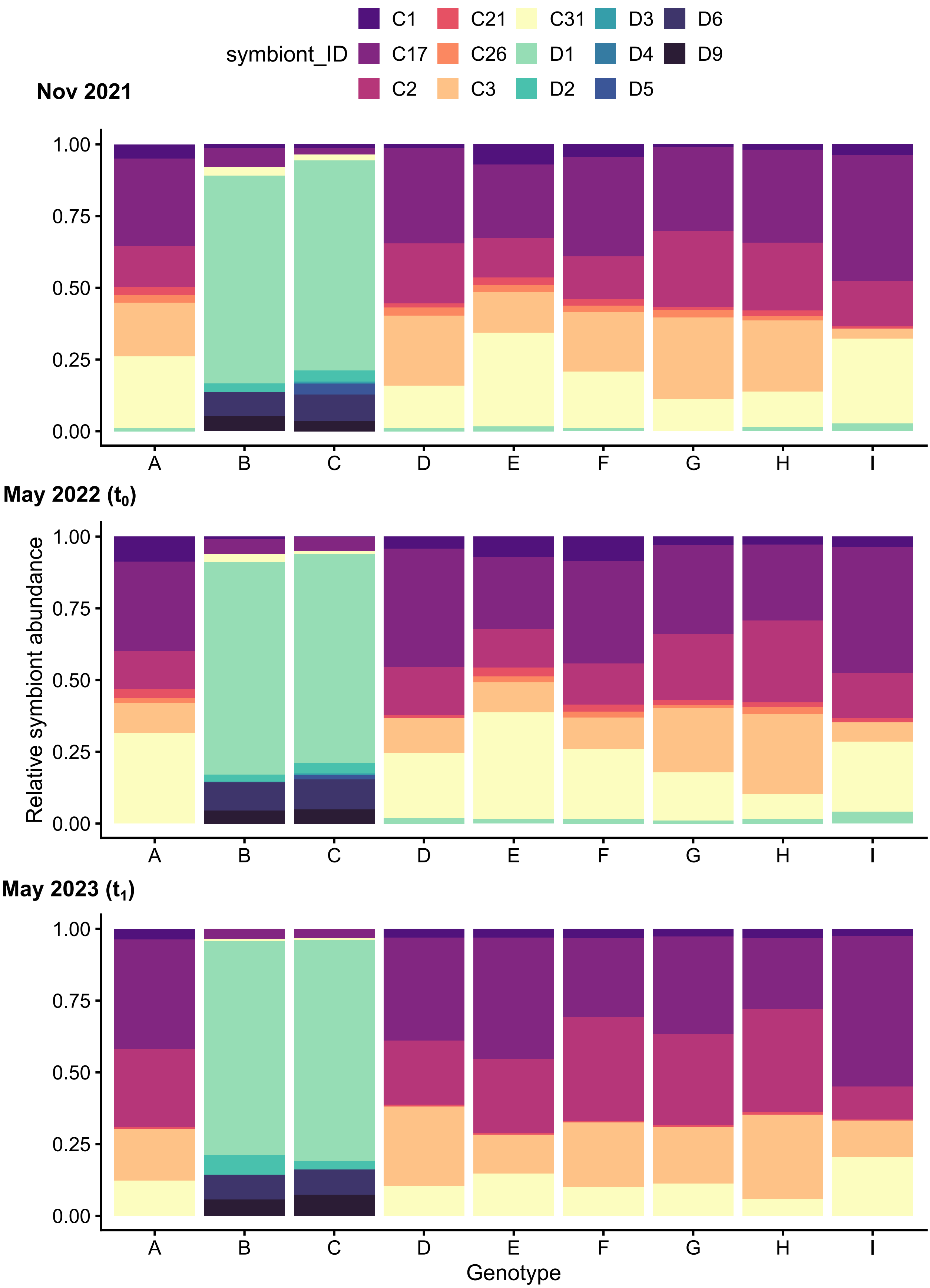


**Fig. S1.** Stacked column plots of symbiont clade prevalence, with means of relative abundances from samples grouped by genotypes for each collection time-point.

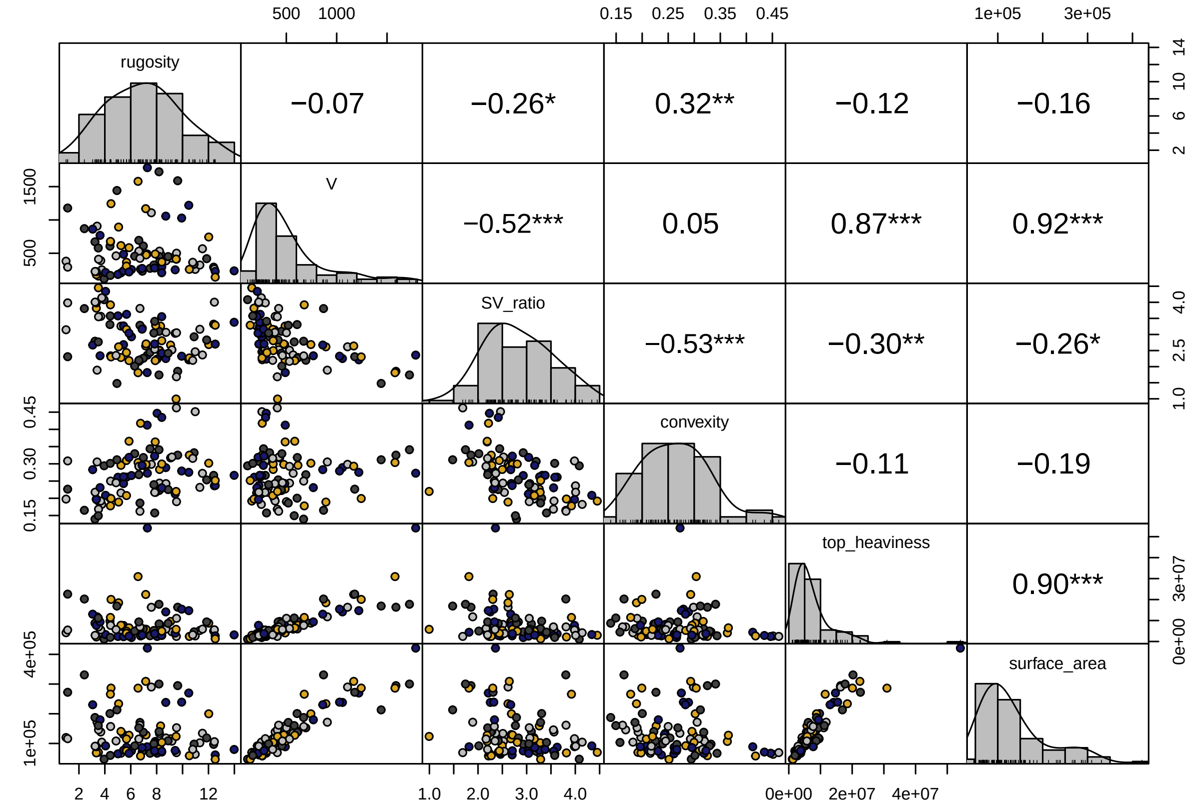


**Fig. S2.** Autocorrelation of untransformed variables. Values above distribution curves indicate the $R^2$ of the linear regression between pairs of variables, and asterisks denote significance levels ($* \leq 0.05; ** \leq 0.01; *** \leq 0.001; **** \leq 0.0001$). Scatterplots below the distributions illustrate the relationships between pairs of variables. Yellow dots = $C_1$ group, dark blue dots = $S_1$ group, light grey dots = $C_0$ group, dark grey dots = $S_0$ group.

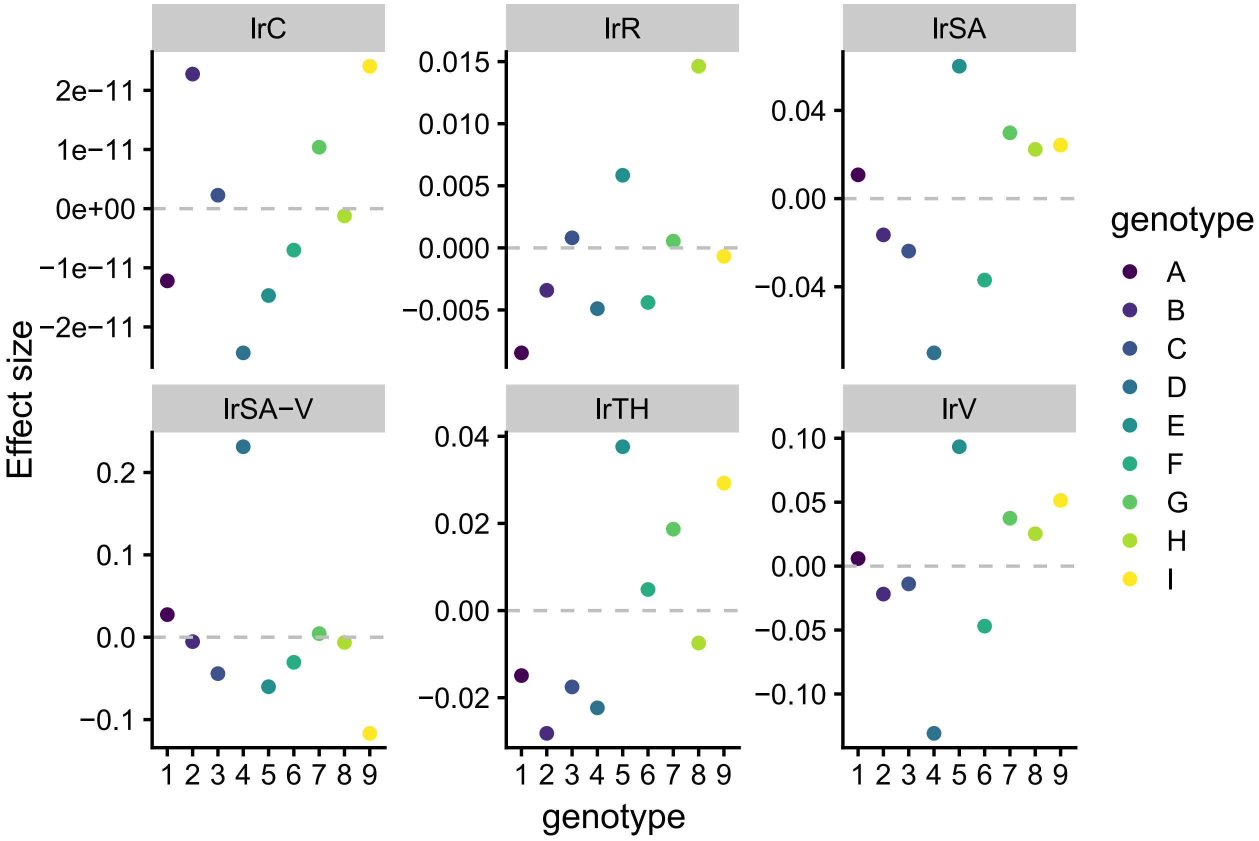


**Fig. S3.** Random effects in the linear mixed models for each morphological trait.

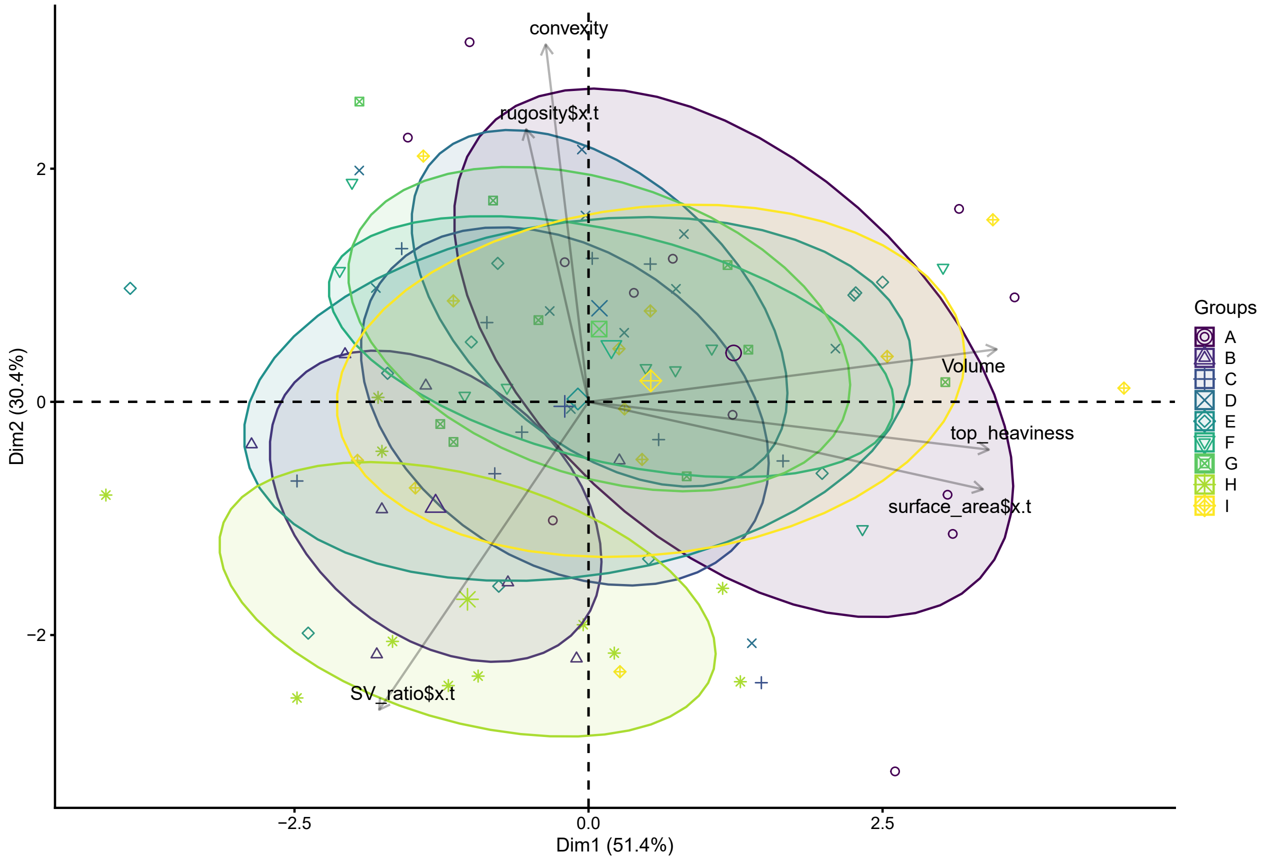


**Fig. S4.** Principal components analysis (PCA) of fragments grouped by genotype. Ellipses show 95% confidence intervals.

**Table S5.** Post-hoc PERMANOVA tests across pairs of genotypes in our dataset. Significance levels: $ns > 0.05; * \leq 0.05; ** \leq 0.01; *** \leq 0.001; **** \leq 0.0001$.

| group1 | group2 | $R^2$ | F value | df1 | df2 | p-value | signif |
|---|---|---|---|---|---|---|---|
| B | E | 0.180 | 3.726 | 1 | 17 | 0.02159784 | * |
| B | F | 0.328 | 7.318 | 1 | 15 | 0.00009999 | **** |
| B | A | 0.294 | 7.479 | 1 | 18 | 0.00009999 | **** |
| B | I | 0.235 | 5.53 | 1 | 18 | 0.00559944 | ** |
| B | H | 0.040 | 0.710 | 1 | 17 | 0.54214579 | ns |
| B | D | 0.314 | 7.797 | 1 | 17 | 0.00049995 | *** |
| B | G | 0.360 | 8.451 | 1 | 15 | 0.00019998 | *** |
| B | C | 0.197 | 3.929 | 1 | 16 | 0.01129887 | * |
| E | F | 0.051 | 0.974 | 1 | 18 | 0.38416158 | ns |
| E | A | 0.106 | 2.498 | 1 | 21 | 0.07769223 | ns |
| E | I | 0.036 | 0.782 | 1 | 21 | 0.47785221 | ns |
| E | H | 0.238 | 6.246 | 1 | 20 | 0.00229977 | ** |
| E | D | 0.065 | 1.392 | 1 | 20 | 0.25347465 | ns |
| E | G | 0.026 | 0.483 | 1 | 18 | 0.67843216 | ns |
| E | C | 0.058 | 1.168 | 1 | 19 | 0.33086691 | ns |
| F | A | 0.090 | 1.885 | 1 | 19 | 0.14918508 | ns |
| F | I | 0.066 | 1.348 | 1 | 19 | 0.25497450 | ns |
| F | H | 0.419 | 12.955 | 1 | 18 | 0.00009999 | **** |
| F | D | 0.004 | 0.078 | 1 | 18 | 0.97550245 | ns |
| F | G | 0.057 | 0.960 | 1 | 16 | 0.38806119 | ns |
| F | C | 0.074 | 1.355 | 1 | 17 | 0.26447355 | ns |
| A | I | 0.033 | 0.758 | 1 | 22 | 0.50144986 | ns |
| A | H | 0.366 | 12.126 | 1 | 21 | 0.00009999 | **** |
| A | D | 0.072 | 1.626 | 1 | 21 | 0.18838116 | ns |
| A | G | 0.148 | 3.291 | 1 | 19 | 0.03519648 | * |
| A | C | 0.086 | 1.883 | 1 | 20 | 0.14178582 | ns |
| I | H | 0.303 | 9.108 | 1 | 21 | 0.00039996 | *** |
| I | D | 0.062 | 1.380 | 1 | 21 | 0.25177482 | ns |
| I | G | 0.085 | 1.764 | 1 | 19 | 0.15428457 | ns |
| I | C | 0.045 | 0.952 | 1 | 20 | 0.40725927 | ns |
| H | D | 0.411 | 13.933 | 1 | 20 | 0.00009999 | **** |
| H | G | 0.423 | 13.193 | 1 | 18 | 0.00009999 | **** |
| H | C | 0.302 | 8.231 | 1 | 19 | 0.00039996 | **** |
| D | G | 0.072 | 1.405 | 1 | 18 | 0.24587541 | ns |
| D | C | 0.065 | 1.329 | 1 | 19 | 0.27397260 | ns |
| G | C | 0.156 | 3.144 | 1 | 17 | 0.02839716 | * |

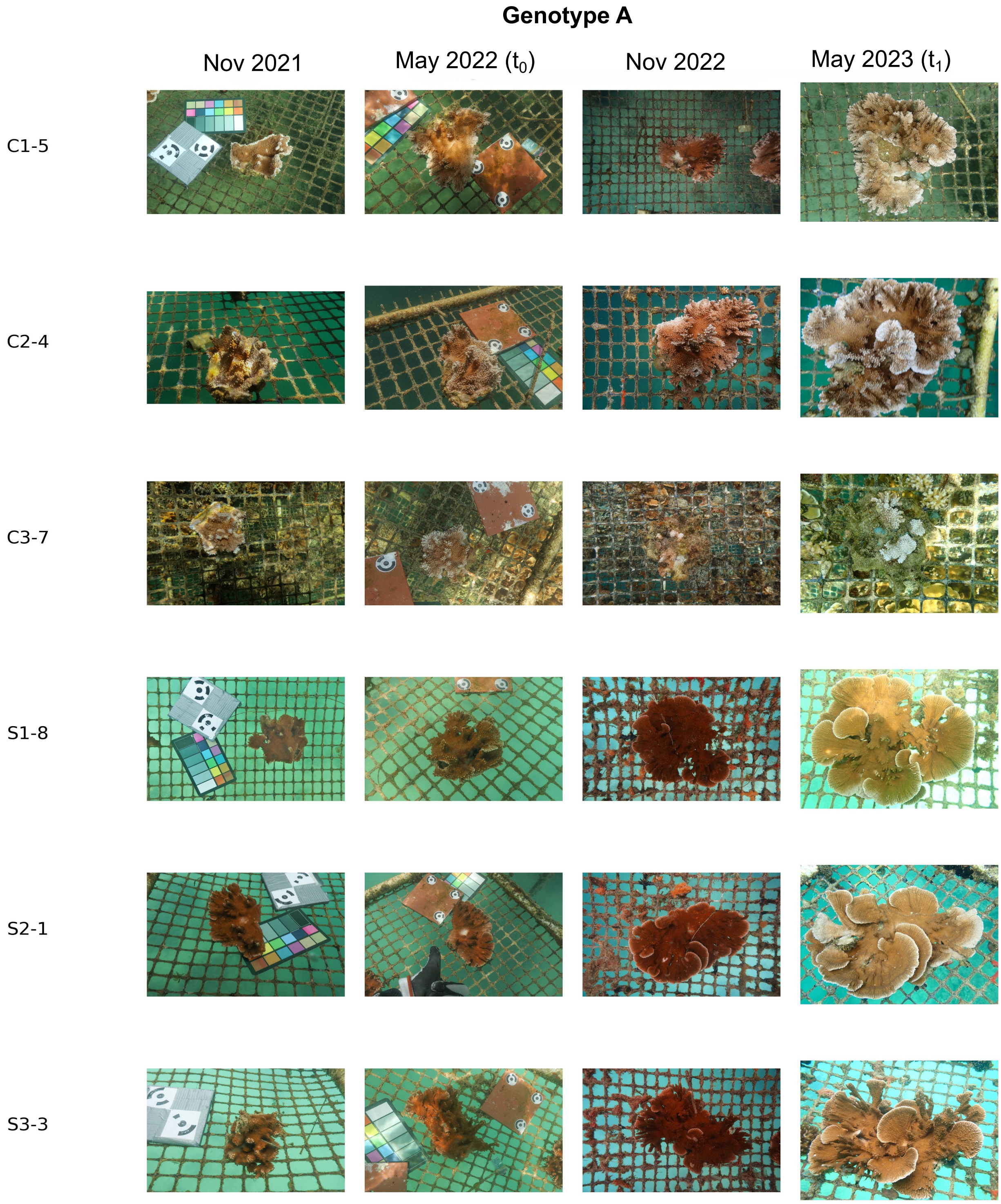


**Fig. S5.** Time-series photographs of control (top three rows) and shaded (bottom three rows) fragments originating from genotype A. The characters on the left correspond to the ID of each coral fragment. Pictures were taken at initial fragmentation (November 2021, first column), $t_0$ (May 2022, second column), six months after $t_0$ (November 2022, third column), and at $t_1$ (May 2023, fourth column). The meshes in the background display square patterns that are approximately 4x4 $cm^2$.

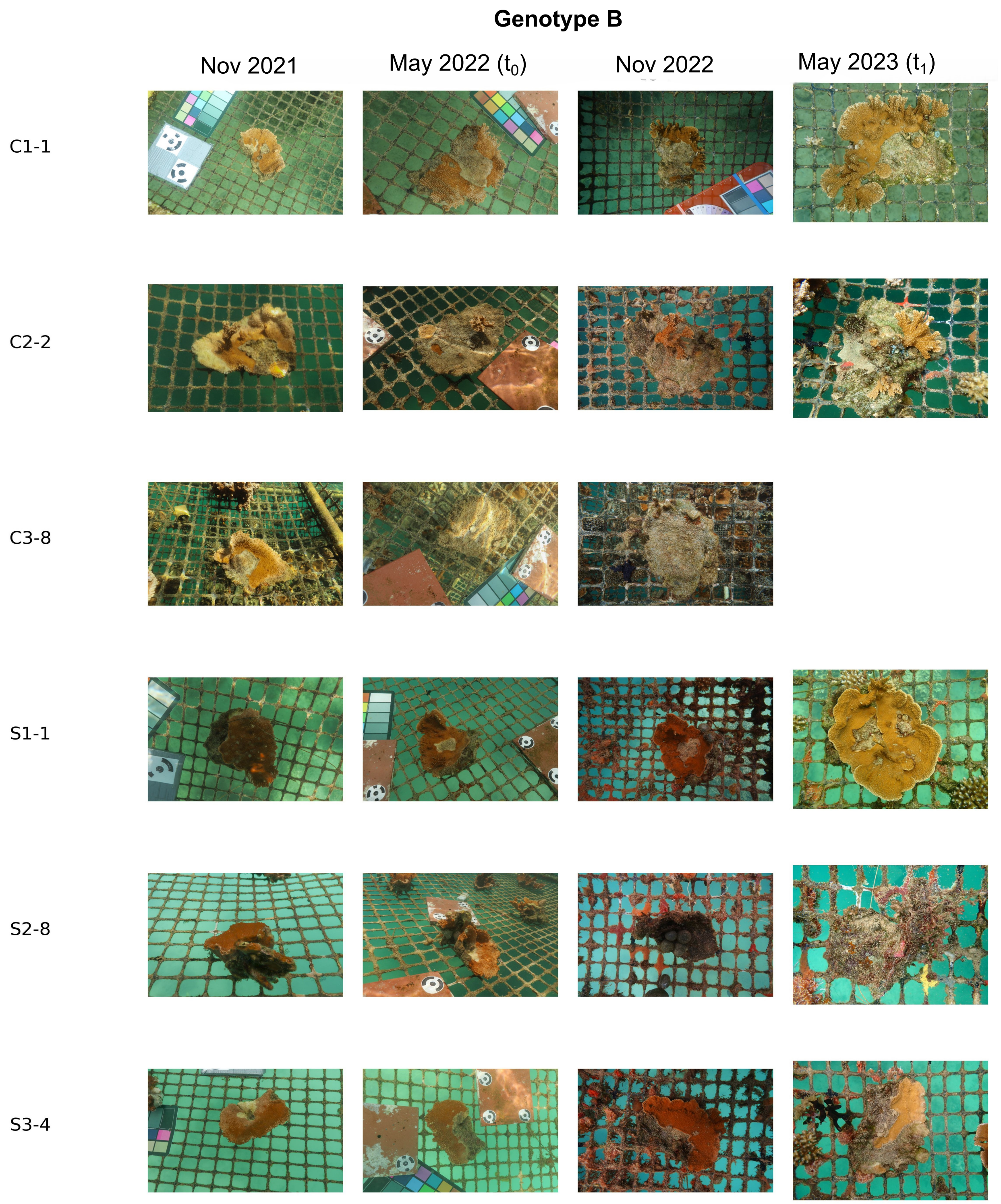


**Fig. S6.** Time-series photographs of control (top three rows) and shaded (bottom three rows) fragments originating from genotype B. The characters on the left correspond to the ID of each coral fragment. Pictures were taken at initial fragmentation (November 2021, first column), $t_0$ (May 2022, second column), six months after $t_0$ (November 2022, third column), and at $t_1$ (May 2023, fourth column). C3-8 was not photographed at $t_1$ as it was lost. The meshes in the background display square patterns that are approximately 4x4 $cm^2$.

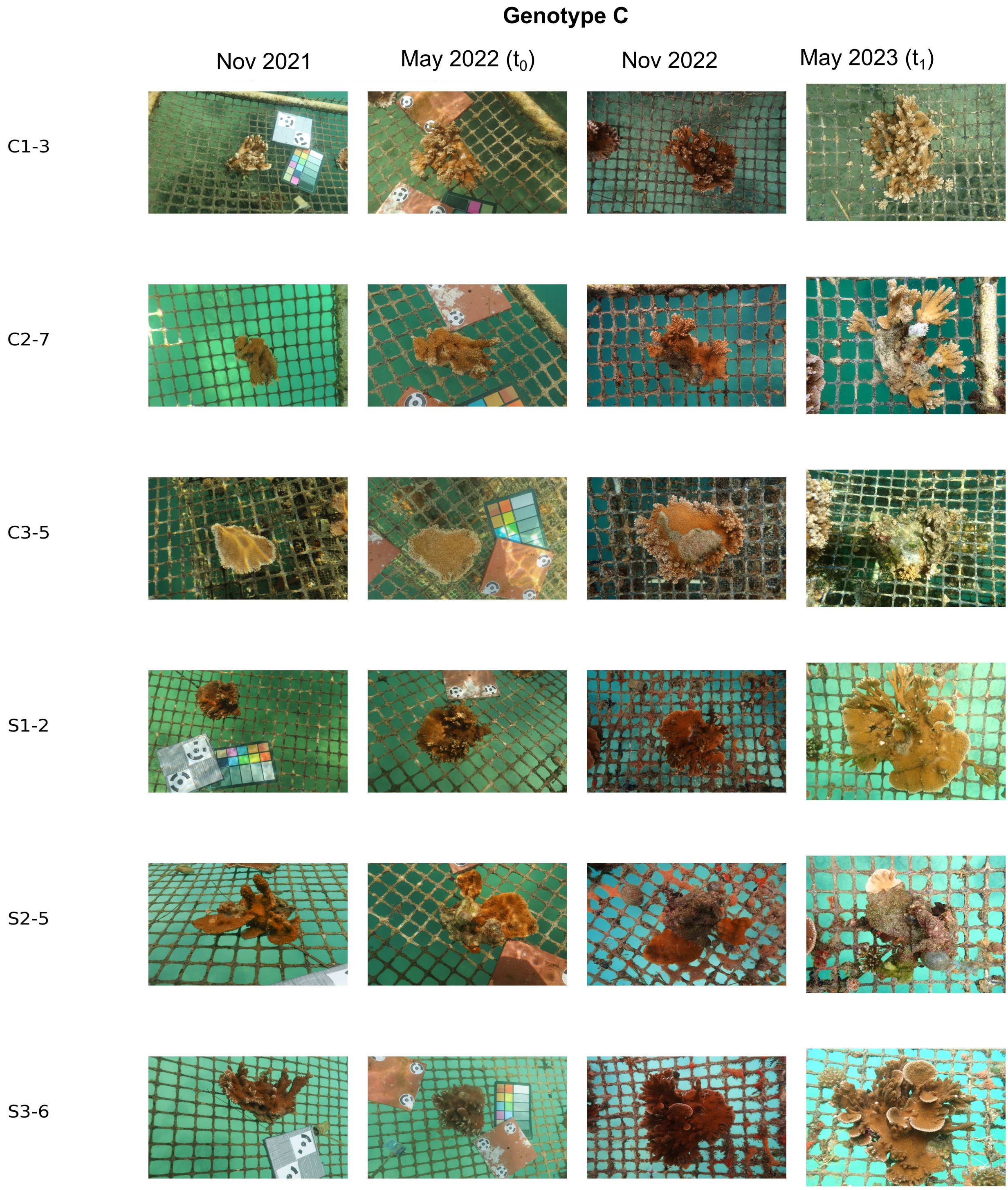


**Fig. S7.** Time-series photographs of control (top three rows) and shaded (bottom three rows) fragments originating from genotype C. The characters on the left correspond to the ID of each coral fragment. Pictures were taken at initial fragmentation (November 2021, first column), $t_0$ (May 2022, second column), six months after $t_0$ (November 2022, third column), and at $t_1$ (May 2023, fourth column). The meshes in the background display square patterns that are approximately 4x4 cm$^2$.

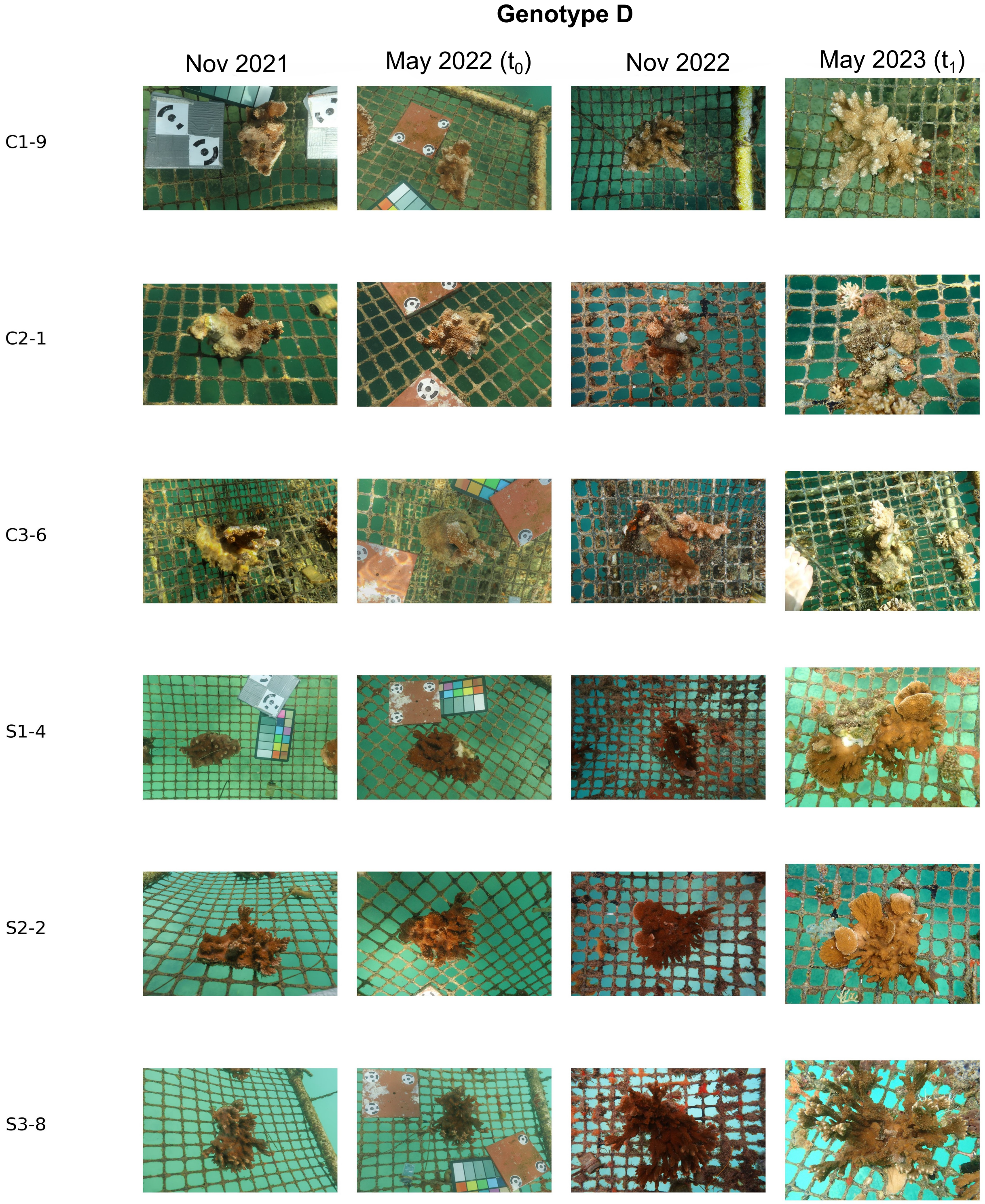


**Fig. S8.** Time-series photographs of control (top three rows) and shaded (bottom three rows) fragments originating from genotype D. The characters on the left correspond to the ID of each coral fragment. Pictures were taken at initial fragmentation (November 2021, first column), $t_0$ (May 2022, second column), six months after $t_0$ (November 2022, third column), and at $t_1$ (May 2023, fourth column). The meshes in the background display square patterns that are approximately 4x4 cm$^2$.

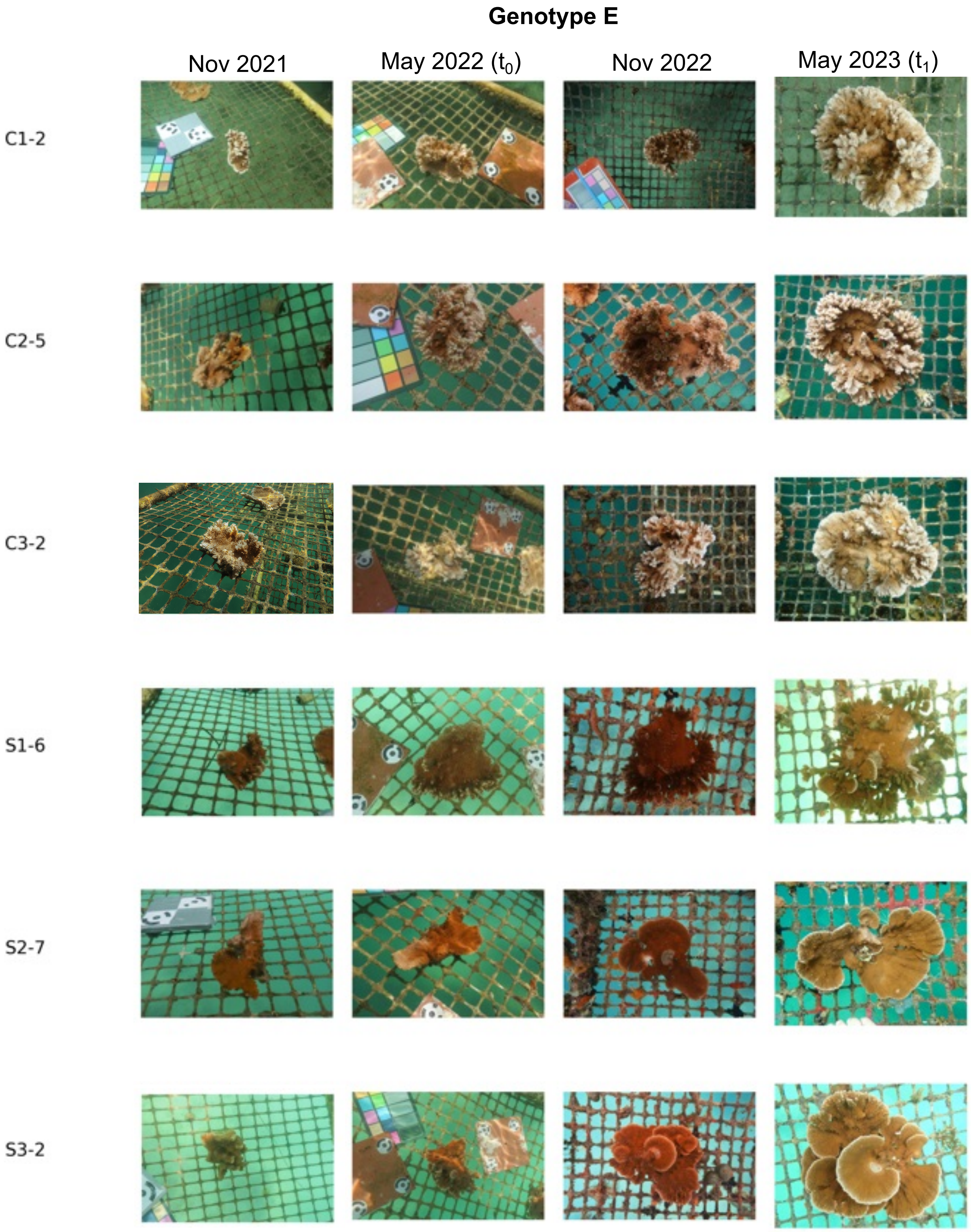


**Fig. S9.** Time-series photographs of control (top three rows) and shaded (bottom three rows) fragments originating from genotype E. The characters on the left correspond to the ID of each coral fragment. Pictures were taken at initial fragmentation (November 2021, first column), $t_0$ (May 2022, second column), six months after $t_0$ (November 2022, third column), and at $t_1$ (May 2023, fourth column). The meshes in the background display square patterns that are approximately 4x4 $cm^2$.

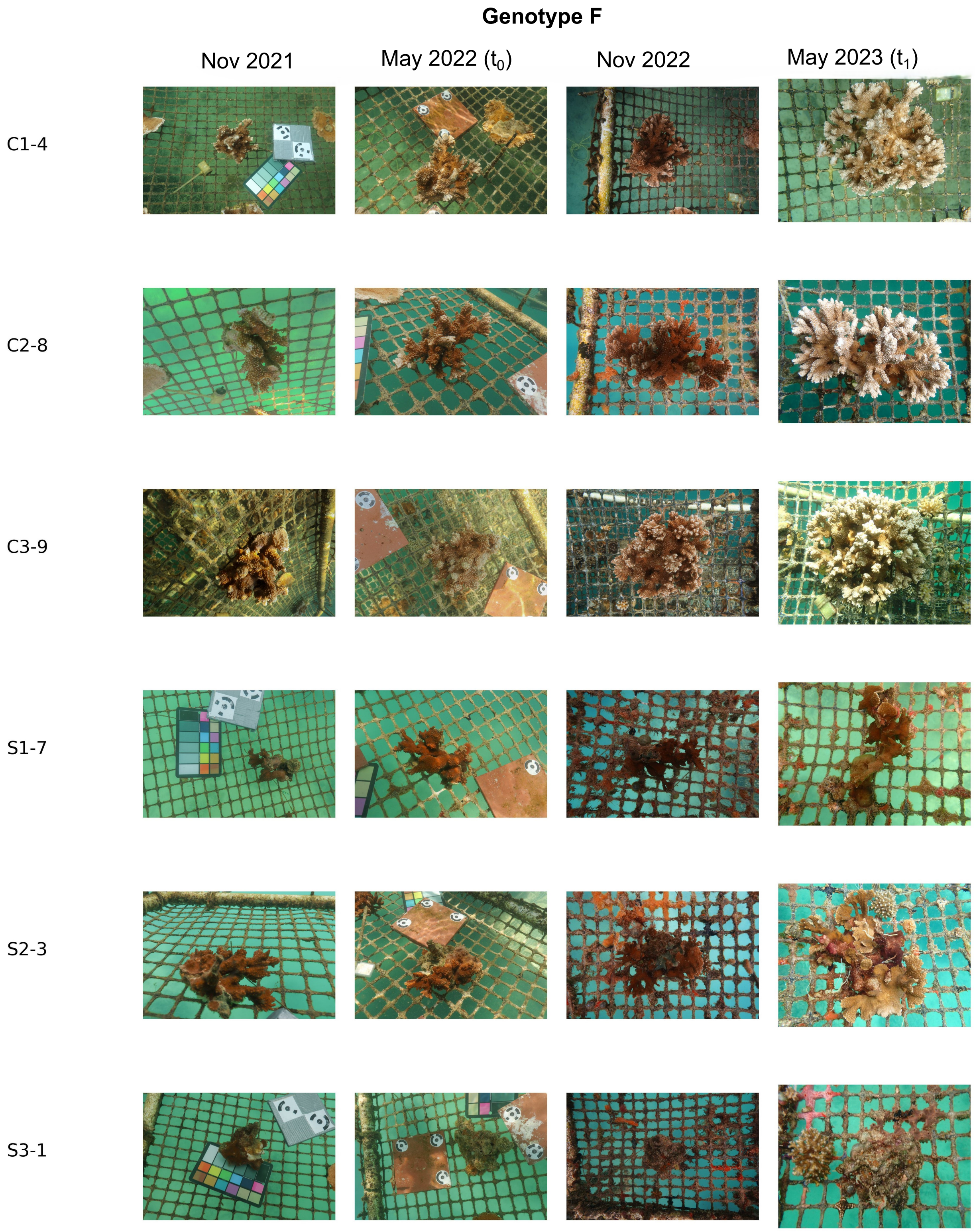


**Fig. S10.** Time-series photographs of control (top three rows) and shaded (bottom three rows) fragments originating from genotype F. The characters on the left correspond to the ID of each coral fragment. Pictures were taken at initial fragmentation (November 2021, first column), $t_0$ (May 2022, second column), six months after $t_0$ (November 2022, third column), and at $t_1$ (May 2023, fourth column). The meshes in the background display square patterns that are approximately 4x4 $cm^2$.

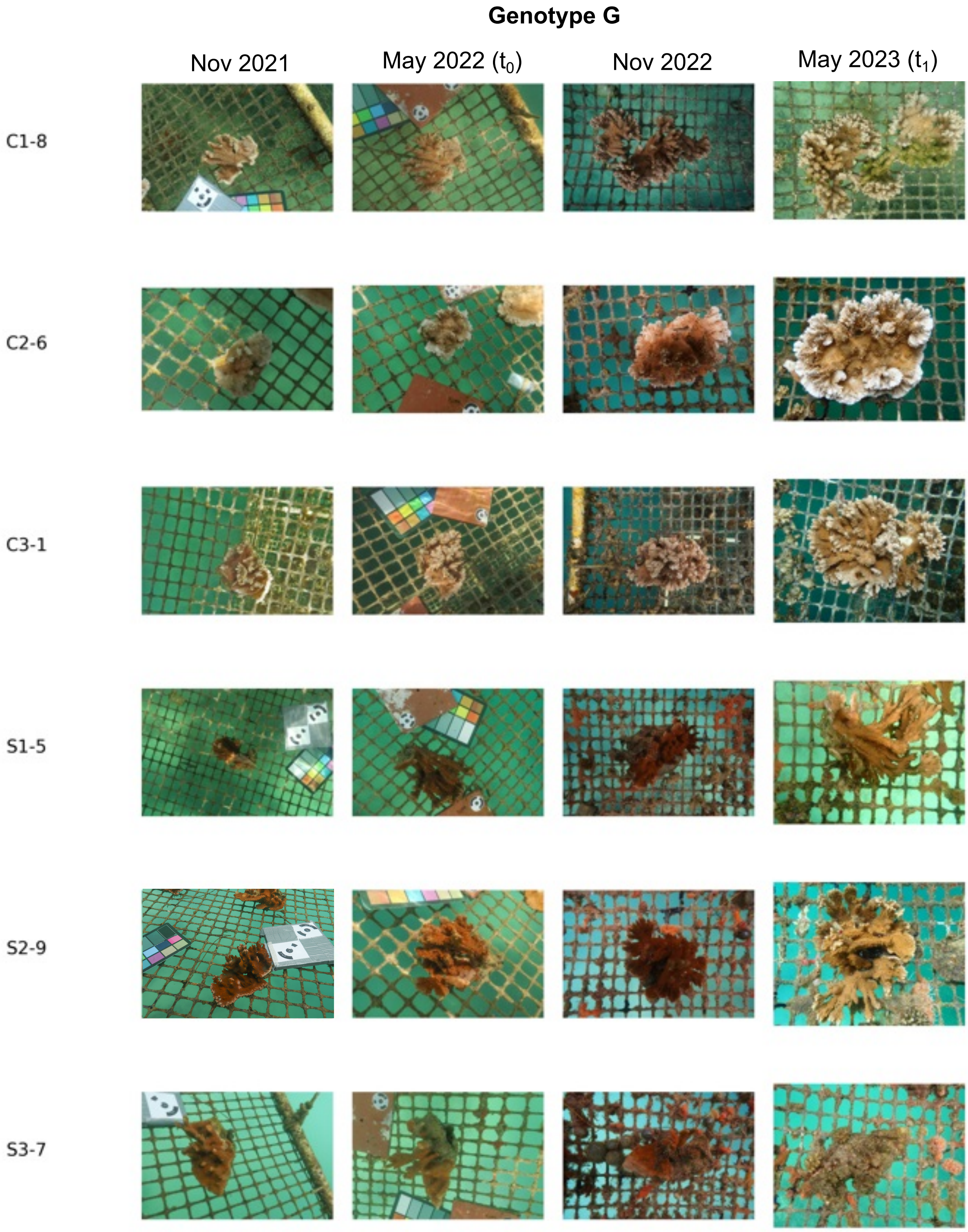


**Fig. S11.** Time-series photographs of control (top three rows) and shaded (bottom three rows) fragments originating from genotype G. The characters on the left correspond to the ID of each coral fragment. Pictures were taken at initial fragmentation (November 2021, first column), $t_0$ (May 2022, second column), six months after $t_0$ (November 2022, third column), and at $t_1$ (May 2023, fourth column). The meshes in the background display square patterns that are approximately 4x4 $cm^2$.

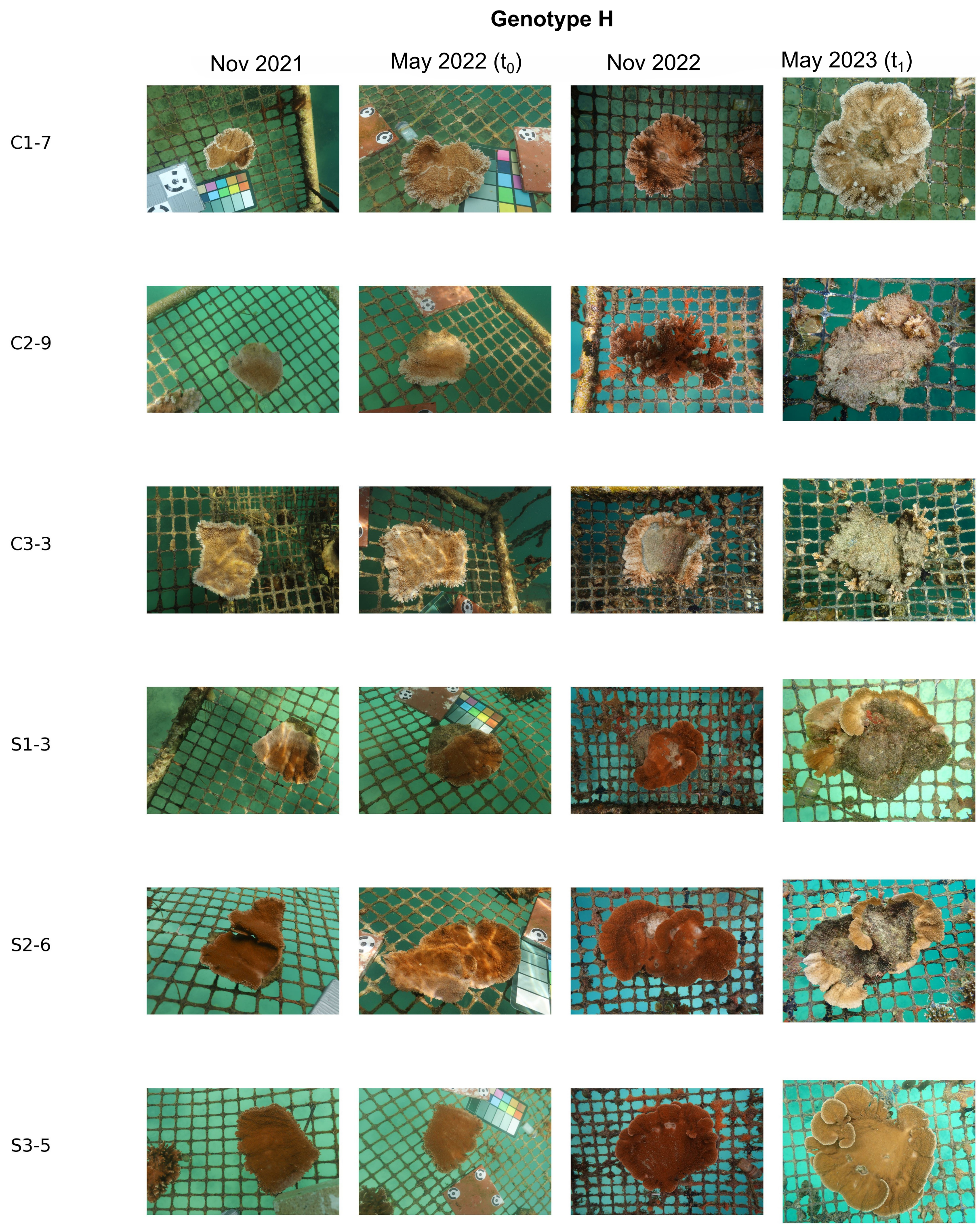


**Fig. S12.** Time-series photographs of control (top three rows) and shaded (bottom three rows) fragments originating from genotype H. The characters on the left correspond to the ID of each coral fragment. Pictures were taken at initial fragmentation (November 2021, first column), $t_0$ (May 2022, second column), six months after $t_0$ (November 2022, third column), and at $t_1$ (May 2023, fourth column). The meshes in the background display square patterns that are approximately 4x4 $cm^2$.

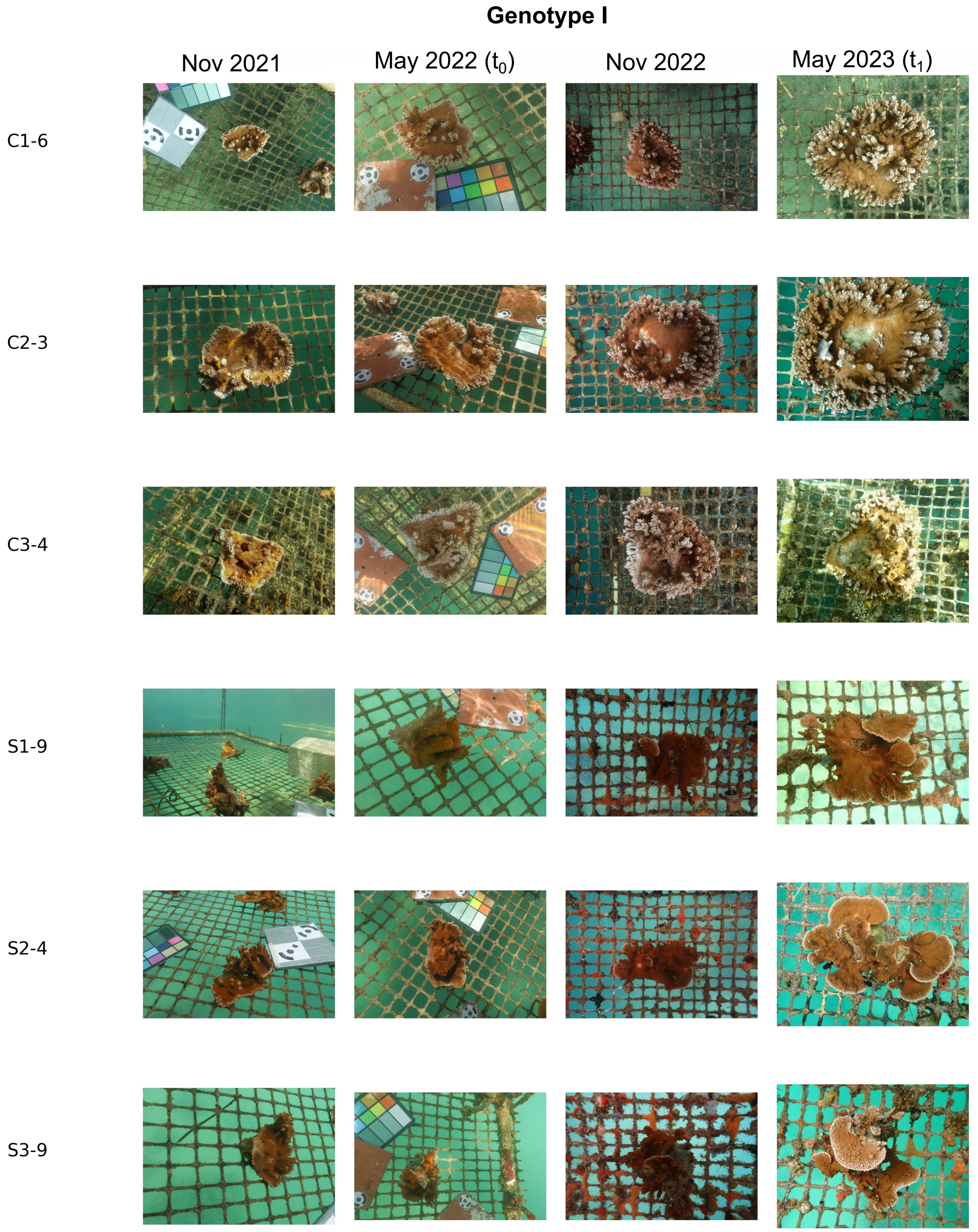


**Fig. S13.** Time-series photographs of control (top three rows) and shaded (bottom three rows) fragments originating from genotype I. The characters on the left correspond to the ID of each coral fragment. Pictures were taken at initial fragmentation (November 2021, first column), $t_0$ (May 2022, second column), six months after $t_0$ (November 2022, third column), and at $t_1$ (May 2023, fourth column). The meshes in the background display square patterns that are approximately 4x4 $cm^2$.
